# Genomic resolution of cryptic species diversity in chipmunks

**DOI:** 10.1101/2022.02.28.482304

**Authors:** Nathanael D. Herrera, Kayce C. Bell, Colin M. Callahan, Erin Nordquist, Brice A. J. Sarver, Jack Sullivan, John R. Demboski, Jeffrey M. Good

**Affiliations:** Division of Biological Sciences, University of Montana, Missoula, Montana, USA; Natural History Museum of Los Angeles County, Los Angeles, California, USA; Department of Biological Sciences, University of Idaho, Moscow, Idaho, USA; Institute for Bioinformatics and Evolutionary Studies (IBEST), University of Idaho, Moscow, Idaho, USA; Department of Zoology, Denver Museum of Nature & Sciences, Denver, Colorado, USA; Wildlife Biology Program, University of Montana, Missoula, Montana, USA

**Keywords:** speciation, hybridization, introgression, phylogenomics, *Tamias*, *Neotamias*

## Abstract

Discovery of cryptic species is essential to understanding the process of speciation and assessing the impacts of anthropogenic stressors. Here, we used genomic data to test for cryptic species diversity within an ecologically well-known radiation of North American rodents, western chipmunks (*Tamias*). We assembled a *de novo* reference genome for a single species (*Tamias minimus*) combined with new and published targeted sequence-capture data for 21,551 autosomal and 493 X-linked loci sampled from 121 individuals spanning 22 species. We identified at least two cryptic lineages corresponding with an isolated subspecies of least chipmunk (*T. minimus grisescens*) and with a restricted subspecies of the yellow-pine chipmunk (*T. amoenus cratericus*) known only from around the extensive Craters of the Moon lava flow. Additional population-level sequence data revealed that the so-called Crater chipmunk is a distinct species that is abundant throughout the coniferous forests of southern Idaho. This cryptic lineage does not appear to be most closely related to the ecologically and phenotypically similar yellow-pine chipmunk but does show evidence for recurrent hybridization with this and other species.

## INTRODUCTION

Species are fundamental units of biodiversity, yet the basic task of species discovery often remains incomplete, even in well-studied taxonomic groups. Whereas some of these difficulties stem from philosophical disagreements on species delimitation (de Queiroz 2007), a far more important source of uncertainty involves the misidentification of cryptic lineages that are phenotypically or ecologically similar to known species (Bickford et al. 2007; Struck et al. 2018). Identifying cryptic diversity is fundamental to understanding both the process of speciation and the conservation of species. The emerging threats of accelerated climate change and habitat loss have only increased the urgency of discovering and more fully accounting for patterns of global biodiversity (Bickford et al. 2007; Delic et al. 2017).

Species delimitation has historically relied on morphological differences. However, molecular data provide evidence that morphological divergence does not always coincide with reproductive isolation. Moreover, the evolutionary history of speciation can be further obscured by hybridization and introgression (Lamichhaney et al. 2017; Fišer et al. 2018). Over the past decade, both theoretical and methodological approaches for genetically based species delimitation have seen considerable progress (Yang and Rannala 2010; Fujita et al. 2012; Edwards and Knowles 2014; Mirarab et al. 2014a; Yang 2015; Mirarab et al. 2016; Luo et al. 2018). In conjunction, advances in high-throughput DNA sequencing provide the ability to reconstruct complex evolutionary histories using sophisticated statistical and phylogenetic approaches that can detect and quantify the extent of gene flow during speciation (Green et al. 2010; Blischak et al. 2018).

Chipmunks of western North America provide a rich system to study speciation and species delimitation across a rapid diversification characterized by ecological adaptation, phenotypic convergence, and overlapping geographic distributions. Of the 25 or 26 (see Burgin et al. 2018) currently recognized chipmunk species (Sciuridae: *Tamias*; but see Patterson and Norris 2016), two are widely distributed lineages occupying central and eastern Asia (*T. sibiricus*) or eastern North America (*T. striatus*; Hall 1981). The remaining 23 species (subgenus *Neotamias*) comprise a recent radiation of habitat specialists and generalists occupying the diverse ecosystems of western North America (Heller 1971; Patterson 1981; Sullivan et al. 2014). Specialization among chipmunks has resulted in strong niche partitioning by habitat and competitive exclusion among co-distributed species (Brown 1971; Heller 1971). Although ecological associations broadly tend to track species boundaries, these associations can break down in sympatry, with chipmunks that occupy similar or transitory habitats showing phenotypic convergence (e.g., in size and pelage coloration) between non-sister species. As a consequence, internal genital bones often provide the only morphological character that is strongly diagnostic of species boundaries (White 1953). Divergence in genitalia is most pronounced in the male genital bone, the baculum (*os penis*), which evolves rapidly among chipmunk species (Callahan 1977; Sutton and Patterson 2000; Sullivan et al. 2014) likely due to strong sexual selection (Eberhard 1985; Simmons 2014). Bacular divergence was long thought to underlie strong reproductive barriers between chipmunk species (Patterson and Thaeler 1982). However, more recent work has shown that some hybridization occurs between species (see Sullivan et al. 2014 for review). In most cases, levels of nuclear introgression have appeared to be relatively low overall (but see Ji et al. 2021), with the persistence of genetically well-defined species boundaries that largely correspond with established taxonomy based on ecological associations and diagnostic genitalia (Hird and Sullivan 2009; Hird et al. 2010; Good et al. 2015; Bi et al. 2019; Sarver et al. 2021).

Genetic studies have also suggested several potential cases of cryptic diversity in this radiation (Demboski and Sullivan 2003; Reid et al. 2012; Sullivan et al. 2014) and the phylogeny of western chipmunks remains unresolved. One potential source for cryptic diversity lies within the widespread least chipmunk (*T. minimus*). For example, *T. m. grisescens* (the Coulee chipmunk, hereafter *grisescens*) is a small, light gray subspecies of *T. minimus* that occupies a restricted range from the Channeled Scablands of central Washington that was identified by Reid and colleagues (2012) as a potential cryptic lineage based on four nuclear loci and mitochondrial DNA (mtDNA).

Another potential example of cryptic diversity occurs within the yellow-pine chipmunk (*T. amoenus*), a widespread species associated with xeric forests throughout western North America. Blossom (1937) described a subspecies of *Eutamias amoenus* (i.e., the Crater chipmunk, *T. amoenus cratericus*, hereafter *cratericus*) as occurring on the periphery of a series of relatively recent lava flows (~10 kya) from Craters of the Moon National Monument and surrounding area of central Idaho. This darker pelage variant of the yellow-pine chipmunk was assumed to reflect local adaptation on the black lava flows of southern Idaho, with rapid transition to more brightly colored *T. amoenus* morphs found in adjacent xeric forest habitats of the region (Figure 1). However, comparison of bacula by White (1953) and Sutton (1982, including the baubellum, the female genital bone; *os clitoris*), of multiple *Tamias* species concluded that *cratericus* may be more closely related to other chipmunk species (*T. umbrinus* or *T. ruficaudus*), based on similar genital morphology and nearby geographic proximity. Finally, the mitochondrial genome sequenced from *cratericus* appears to be fairly divergent from other species, and has been detected at an additional locality to the north of Craters of the Moon (Demboski and Sullivan 2003) suggesting that *cratericus* may be more widespread (Reid et al. 2012).

**Figure 1.**
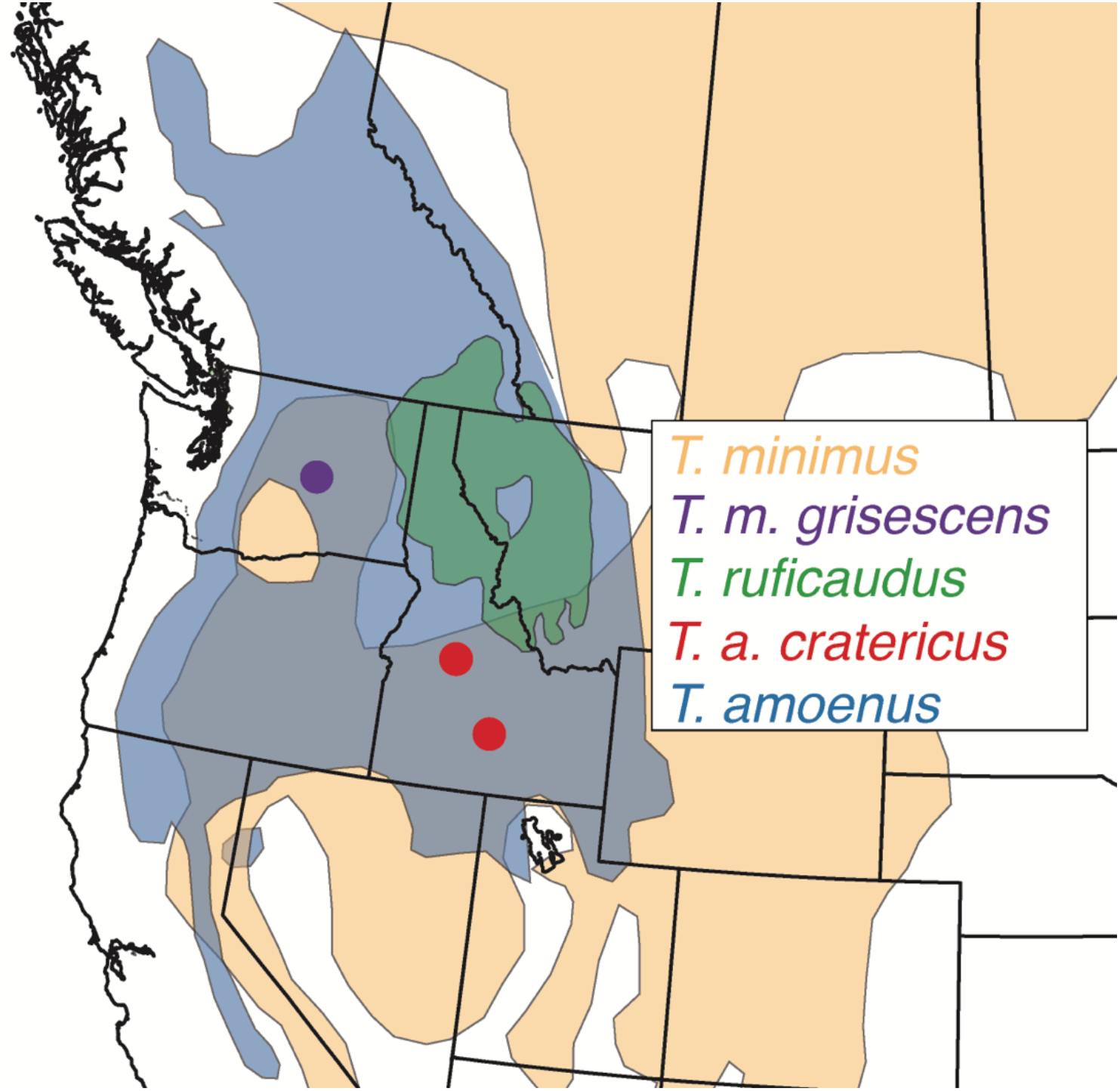
Two potential cryptic species of chipmunk. *T. m. grisescens* (Purple) occupies a restricted range from the Channeled Scablands of central Washington. *T. a. cratericus* (Red) is described as a locally adapted form of the yellow-pine chipmunk (*T. amoenus*).

Here, we use linked-read sequencing to generate a reference genome for the least chipmunk (*T. minimus*). We then use new and published targeted capture sequencing data from 21,551 autosomal and 493 X-linked loci, and complete mitochondrial genomes to infer the phylogenetic relationships among 22 described chipmunk species, including the enigmatic *cratericus* and *grisescens* chipmunks. We then use additional population-level sequencing to characterize the geographic range and evolutionary history of *cratericus* relative to three other potentially co-distributed chipmunk species.

## MATERIALS AND METHODS

### SAMPLING AND ETHICS STATEMENT

To assess genus-wide diversity, we combined extensive fieldwork with collections from several natural history museums to obtain samples from 14 western chipmunk species and the eastern chipmunk, *Tamias striatus*, as an outgroup (Supplementary Table S1). We combined data from these samples with published data (Bi et al. 2019; Sarver et al. 2021) from seven additional species of western chipmunks resulting in broad sampling covering 22 of the 26 described species of chipmunk (57 individuals). Not included were samples of *T. solivagus*, *T. durangae*, and *T. bulleri* (all western chipmunk species endemic to Mexico), and the Eurasian *T. sibiricus*. To better characterize the geographic and ecological extent of *cratericus*, we collected an additional 64 individuals from 10 populations throughout central Idaho covering a broad range of habitats from the lava flows of Craters of the Moon up to the subalpine zone. All animals sampled for this project were collected following procedures approved by the University of Montana Animal Care and Use Committee (protocol numbers AUP 042-16JGDBS & 022-10JGDBS). We used acceptable field collecting methods, conducted according to the recommendations of the American Society of Mammalogists Animal Care and Use Committee (Sikes and Mammalogists 2016).

### GENOME SEQUENCING AND ASSEMBLY

Liver tissue from a female *T. minimus* was frozen in liquid nitrogen immediately after euthanasia and sent to the DNA Technologies and Expression Analysis Core at the UC Davis Genome Center for DNA extraction and library preparation. High molecular weight genomic DNA was isolated following Jain et al. (2018). Briefly, 40 mg of flash frozen liver tissue was homogenized by grinding in liquid nitrogen and lysed with 2 ml of lysis buffer containing 10 mM Tris-HCl pH 8.0, 25 mM EDTA, 0.5% (w/v) SDS and 100μg/ml Proteinase K. The lysate was cleaned with equal volumes of phenol/chloroform using phase lock gels (Quantabio Cat # 2302830). The DNA was precipitated by adding 0.4X volume of 5M ammonium acetate and 3X volume of ice-cold ethanol, washed twice with 70% ethanol, and resuspended in buffer (10mM Tris, pH 8.0). The integrity of the high-molecular-weight genomic DNA was verified on a Pippin Pulse gel electrophoresis system (Sage Sciences, Beverly, MA). Following DNA isolation, a 10X Genomics Chromium library was prepared following the manufacturer’s protocol and sequenced on the Illumina HiSeq X platform by Novogene Corporation. We then used Supernova V2.0.0 (Weisenfeld et al. 2017) to generate a *de novo* genome assembly from the 10X Genomics Chromium library sequence data using default settings with --maxreads=1135000000 (the number of reads estimated to yield 56× coverage for an estimated genome of ~2.3 Gigabases; Supplementary Table S2).

### EXON CAPTURE ENRICHMENT AND SEQUENCING

We generated new genome-wide resequencing data from 112 individuals and 15 chipmunk species (including *T. striatus* as an outgroup; Supplemental Table S1) using a custom in-solution targeted capture experiment designed by Bi et al. (2019) to enrich and sequence exons (including flanking introns and intergenic regions) from 10,107 nuclear protein-coding genes [9.4 Megabases (Mb) total and the complete ~16.5 kilobase (kb) mitochondrial genome]. This capture strategy was iteratively developed from a series of genomic experiments. First, RNA-seq was used to sequence and assemble transcriptomes from multiple tissue types sampled from a single chipmunk species (*T. alpinus*), which served as the reference for exome capture probe design (Bi et al. 2012). These transcriptome data were used to develop an array-based capture targeting ~8,000 nuclear genes as well as the complete mitochondrial genome, resulting in ~6.9 Mb of assembled exonic regions (including flanking introns and intergenic regions; Bi et al. 2012; Bi et al. 2013; Good et al. 2015). Finally, an additional ~2.4 Mb of genic regions were identified from the original transcriptome through AmiGO and NCBI protein databases and were added to the capture design. Published data from Sarver et al. (2021; *T. canipes*, *T. cinereicollis*, *T. dorsalis*, *T. quadrivitattus*, *T. rufus*, and *T. umbrinus*) were based on the earlier array-based design and thus represent an overlapping subset of the genomic regions targeted here.

To localize the targeted genomic regions within the *de novo T. minimus* genome, we first used BLAT to identify the best hit in the reference. We then sorted and filtered the contigs to include only the best match hits, discarding contigs with equally likely hits across the genome. To identify X-linked contigs in the genome assembly, we calculated the normalized female versus male ratio of sequencing coverage for targeted regions mapping to each contig, which should be approximately 2:1 for genes on the X chromosome.

Total genomic DNA was isolated from heart or liver tissue using Qiagen DNEasy DNA blood and tissue kits. Illumina sequencing libraries were prepared for each sample following the protocol of Meyer and Kircher (2010). Sequencing libraries were then enriched in solution using probes designed as described above and manufactured by NimbleGen (SeqCap EZ Developer kits). Hybridization reactions were performed in four separate equimolar pools of indexed libraries. To assess capture efficiency, we first sequenced a subset of pooled libraries on an Illumina MiSeq sequencer (150 bp paired-end reads) at the University of Montana genomics core (Missoula, MT). Final pooled libraries were sequenced on an Illumina HiSeq 2500 (150 bp paired-end reads) at the University of Oregon (Eugene, OR).

### ILLUMINA DATA PROCESSING AND GENOTYPING

Raw read data were cleaned using the *HTStream* pipeline (available from https://github.com/s4hts/HTStream; last accessed July 21, 2018), to trim adapters and low-quality bases, merge overlapping reads, and remove putative PCR duplicates. Cleaned reads were mapped to the reference genome using the BWA-MEM alignment algorithm as implemented in BWA v0.7.15 (Li and Durbin 2009; Li 2013) with default options and then sorted with *samtools* v1.4 (Li et al. 2009). Mapped reads were assigned read groups and duplicate reads were identified post-mapping using Picard v2.5.0 (http://broadinstitute.github.io/picard/). Initial capture efficiency statistics were calculated using QC3 v1.33 (Guo et al. 2014). Regions with insertions or deletions (indels) were identified and realigned, and single nucleotide variants (SNVs) were called using UnifiedGenotyper within the Genome Analysis Toolkit (GATK) v3.6 (McKenna et al. 2010; DePristo et al. 2011; Van der Auwera et al. 2013) with the EMIT_ALL_SITES argument set. This resulted in a final VCF with a call at every position. We then soft filtered based on minimum mapping quality (MQ < 20), a minimum quality (QUAL < 20), a minimum quality by depth (QD < 2), and a minimum sequencing depth (DP < 5), resulting in a final filtered single sample VCF file used for all downstream analyses.

### PHYLOGENETIC INFERENCE AND SPECIES DELIMITATION

To obtain sequence alignments for phylogenetic analyses, we constructed consensus sequences of the targeted regions using the filtered VCF for each individual by injecting all confidently called sites back into the original reference using FastaAlternateReferenceMaker within the GATK. IUPAC ambiguity codes were inserted at putative heterozygous positions using the - IUPAC flag. Remaining ambiguous positions (MQ < 20, QUAL < 20, QD < 2, DP< 5) were hard masked (i.e., replaced with an “N”) using GNU awk and bedtools v2.25 (Quinlan and Hall 2010).

We used a two-tiered approach to infer phylogenetic relationships within *Tamias.* First, we used a concatenated alignment of the targeted regions and 100 base pairs (bp) of flanking sequence to estimate a single bifurcating phylogeny using a maximum likelihood (ML) search as implemented in IQ-Tree v.1.5.5 (Nguyen et al. 2015). For all phylogenetic analyses, we ran separate analyses for autosomal and X-linked regions. We use ModelFinder Plus (-m MFP) (Kalyaanamoorthy et al. 2017) and Bayesian information criterion (BIC) to select the best nucleotide substitution model for the concatenated data. To assess nodal support, we performed 1000 ultrafast bootstrap replicates (UFBoot) with the “-bb” command (Minh et al. 2013; Hoang et al. 2017). Concatenation of multiple loci ignores coalescent stochasticity and can result in erroneously high bootstrap support for nodes that may show substantial incongruence when considered on a per locus basis (Kumar et al. 2012). Therefore, we also calculated gene concordance factors (gCF) and site concordance factors (sCF) as implemented in IQ-Tree v. 2.0.6 (Minh et al. 2020a; Minh et al. 2020b). Gene concordance factors are the proportion of inferred single locus trees (gene trees) that support a particular branch in the reference tree (ML concat tree), whereas site concordance factors are the proportion of sites within the concatenated alignment that support a branch in the reference tree.

To obtain gene trees, we extracted alignments from the targeted regions and 100 bp of flanking sequence by combining all targets found within stepped 50 kb intervals using bedtools v2.2.5 and combined regions by scaffold using AMAS (Borowiec 2016). This approach was chosen to capture the spatial distribution of individual targets while ensuring we include enough informative sites to infer independent phylogenetic histories. Although interval size is somewhat arbitrary here, we considered a range of interval sizes and found smaller intervals (≤ 10 kb) resulted in many intervals without targets and few informative sites. Likewise, much larger interval sizes (≥ 100 kb) were more likely to conflate independent phylogenetic histories. For each interval, we filtered positions with missing data for >85% of individuals using AMAS and excluded intervals containing less than 1 kb of capture data. We then used IQ-Tree v.1.5.5 to estimate local maximum likelihood trees with 1000 ultra-fast bootstraps (using -m MFP, -bb 1000), with *T. striatus* as the outgroup. Gene trees, when necessary, were unrooted using the R package *ape* (Paradis et al. 2004). We then used unrooted gene trees to estimate a consensus species tree using ASTRAL-III v5.6.3 (Zhang et al. 2018). Assuming sets of independent and accurately estimated gene trees, ASTRAL estimates an unrooted species tree by finding the species tree that has the maximum number of shared induced quartet trees with the given set of gene trees (Mirarab et al. 2014a; Mirarab et al. 2014b).

Next, we estimated coalescent-based species trees with SVDquartets (Chifman and Kubatko 2014) using 100,000 random quartets and 100 bootstrap replicates as implemented in PAUP* v4.0a166 (Swofford 2003). SVDquartets assesses support for quartets of taxa directly from site-pattern frequencies of variable sites only. This approach differs from summary methods, such as ASTRAL, because it does not independently estimate gene trees, avoiding the issue of gene-tree estimation error (Chifman and Kubatko 2015; Chou et al. 2015). For SVDquartets, we extracted single nucleotide variants (SNVs) distanced at least 10 kb along the genome and excluded sites with missing information for >85% of the individuals. For both SVDquartets and ASTRAL analyses, species trees were estimated with and without assigning species identities and using sites or intervals for autosomal and X-linked loci separately as previously described. When applicable, we included the eastern chipmunk, *T. striatus*, as the outgroup.

Finally, we estimated a species tree using the Bayesian program BPP version 4.4.1 (Rannala and Yang 2017; Flouri et al. 2018). BPP uses a full likelihood-based implementation of the multispecies coalescent and accommodates uncertainty due to gene tree heterogeneity (both the topology and branch lengths) and incomplete lineage sorting. We first performed preliminary runs using a diffuse prior (shape parameter α=3) to assess the fit of the prior means with our data. For our model parameters, we assigned inverse-gamma priors to the population size parameter θ [∼IG (3, 0.002) with mean 0.00], and to the divergence times τ [∼ IG (3, 0.01) with mean 0.005 for the age of the root]. We assessed convergence based on consistency across 5 replicate MCMC runs, using a different starting tree and seed for each. For each run, we generated 200,000 samples with a sample frequency of two iterations after a burn-in of 16,000 iterations. While BPP can potentially analyze 1000’s of genetic loci, we found that runtimes were excessive even with our relatively modest genome-wide capture dataset. Therefore, we explored several pruning options and settled on selecting the largest contig with a minimum interval distance of 1 Mb. This resulted in a sequence alignment consisting of 88 loci (ranging in size from ~1,000-4,600 bp; median 2,343 bp) for a total alignment length of 209,725 bp.

### INFERENCE OF POPULATION STRUCTURE

We plotted PC1 and PC2 from a principal components analysis (PCA) using the R package SNPRelate (Zheng et al. 2012). To estimate individual co-ancestry, we performed ML estimation of individual admixture proportions using the program ADMIXTURE v.1.30 (Alexander et al. 2009). We tested values of K (the number of population clusters) ranging from 1-6 under the default settings in ADMIXTURE and selected the model with the lowest cross-validation error. For both PCA and population cluster analyses, we used a filtered SNP dataset where we applied LD-based pruning to filter our SNP dataset using PLINK v1.9 (Chang et al. 2015) to minimize non-independence due to linkage. We used a 1 kb sliding window with a step size of 100 bp and pairwise *r*^2^ of 0.8 as a cut-off for removing highly linked SNPs. This resulted in a final set of 93,247 SNPs.

### INTROGRESSION AND PHYLOGENETIC NETWORKS

An advantage of SVDquartets is that it detects deviations in site-pattern frequencies expected under the multi-species coalescent by evaluating support for all three resolutions for each quartet, similar to the ABBA-BABA test (Green et al. 2010; Durand et al. 2011). Therefore, we used the software package HyDe (Blischak et al. 2018) to test for hybridization using the invariants framework of SVDquartets. Here, the quartet with the majority of the support should correspond to the species tree. With incomplete lineage sorting (ILS) and in the absence of introgression, the two minor quartet topologies should show similarly low support. Alternatively, introgression will lead to an imbalance of support towards the topology with the two taxa exchanging alleles as sister taxa (Pease and Hahn 2015; Blischak et al. 2018; Kubatko and Chifman 2019). HyDe also estimates γ, which is the parental contribution in a putatively hybrid genome, where a value of 0.5 would indicate a 50:50 genomic contribution from each parent.

We also modeled hybridization and ILS under the coalescent network framework as implemented in PhyloNet 3.6.6 (Yu and Nakhleh 2015). Networks were computed under Maximum Pseudo-Likelihood (*InferNetwork_MPL*) (Yu and Nakhleh 2015) using rooted autosomal gene trees. We modeled 0 to 5 migration events, associated individuals to species (option -a), and optimized branch lengths and inheritance probabilities to compute likelihoods for each proposed network (option -o). We calculated Akaike information criteria corrected for small sample sizes (AICc) and Bayesian Information Criteria (BIC) using the highest likelihoods per run to compare the resulting networks (Yu et al. 2012; Yu et al. 2014). Networks were visualized with IcyTree (https://icytree.org; last accessed Dec 2020).

## RESULTS

### DRAFT GENOME ASSEMBLY AND SEQUENCE CAPTURE

We generated ~217 Gb of *T. minimus* genomic sequence data using 10X Genomics technology (Pleasanton, CA, USA) and assembled a 2.48 Gb draft genome with a contig N50 of 196.09 kb and as scaffold N50 of 58.28 Mb, respectively (Supplementary Table S2). We then analyzed up to 9.4 Mb of sequence capture data from 121 individuals, combining newly generated (112 individuals from 15 species) and published data (nine individuals from seven species; Bi et al. 2019; Sarver et al. 2021; Supplementary Table S1) for 21 western chipmunk species and the outgroup *T. striatus.* Sequencing efforts produced ~4 million reads per sample with an average of 0.5% of targets showing no coverage. Approximately 75% of raw reads were unique, resulting in an average target coverage of 33× across samples (range: 5–91×) with ~73% of targeted bases sequenced to at least 10× coverage (Supplementary Table S3). Comparison of male versus female coverage resulted in the identification of three contigs (79.7 Mb, 35.6 Mb, 5.9 Mb) in the genome assembly that are likely on the X chromosome.

### THE *TAMIAS* PHYLOGENY

We first estimated a genus-wide ML phylogeny from a concatenated set of 21,551 autosomal (5,365,556 bp) and 493 X-linked (105,671 bp) loci from the combined capture datasets, as well as an ML phylogeny based on the full mtDNA genome (Figure 2A; Supplemental Figure S1-S2). The concatenated autosomal ML analysis recovered a fully resolved phylogeny with high support for all branches (UFBoot > 90%) and is concordant with previous studies (Reid et al. 2012; Sullivan et al. 2014; Sarver et al. 2021). *Tamias m. grisescens* appeared as a distinct lineage in these analyses and was not most closely related to other *T. minimus* samples. Likewise, *cratericus* was a distinct, monophyletic lineage that was most closely related to *T. ruficaudus* and not other *T. amoenus*. Within *cratericus*, we also recovered a deep split between the two sampled localities (hereafter *cratericus* lineages A and B; Figure 2A). ML reconstruction of the mitochondrial genome resulted in a different topology from the nuclear dataset (Supplemental Figure S2) and was largely concordant with previous studies based on the cytochrome *b* gene (Demboski and Sullivan 2003; Good et al. 2003; Good et al. 2008; Reid et al. 2012; Good et al. 2015; Sarver et al. 2017) with *cratericus* most closely related to *T. quadrimaculatus*, and then to *T. minimus*. Our mitochondrial genome reconstruction also recovered an individual from the putative *cratericus* lineage B (CRCM06-160; Supplemental Figure S2) as having a *T. amoenus* mtDNA genome, consistent with introgression. The multispecies-coalescent species trees estimated from ASTRAL (158 genes trees estimated from 50 kb genomic intervals) and SVDquartets (13,482 unlinked SNPs) was largely concordant with the results of the concatenated ML analyses. Strong support was recovered for the branching relationships among species groups within *Tamias* (ASTRAL posterior probabilities > 0.9 and SVDquartets bootstrap support > 90). All analyses recovered both *grisescens* and *cratericus* as distinct, monophyletic lineages. However, one notable difference across analyses was the relationship of *cratericus* with respect to *T. ruficaudus* and other *T. amoenus*. Both concatenated ML analyses and ASTRAL recovered *cratericus* as the sister lineage to *T. ruficaudus*, whereas SVDquartets recovered *cratericus* as more closely related to other *T. amoenus* but with weak support (Supplemental Figures S3-S6). Overall, the splits between *T. amoenus*-*cratericus*-*T. ruficaudus* were not well-supported and remained unresolved (Figure 2A; Supplemental Figure S1-S6).

**Figure 2.**
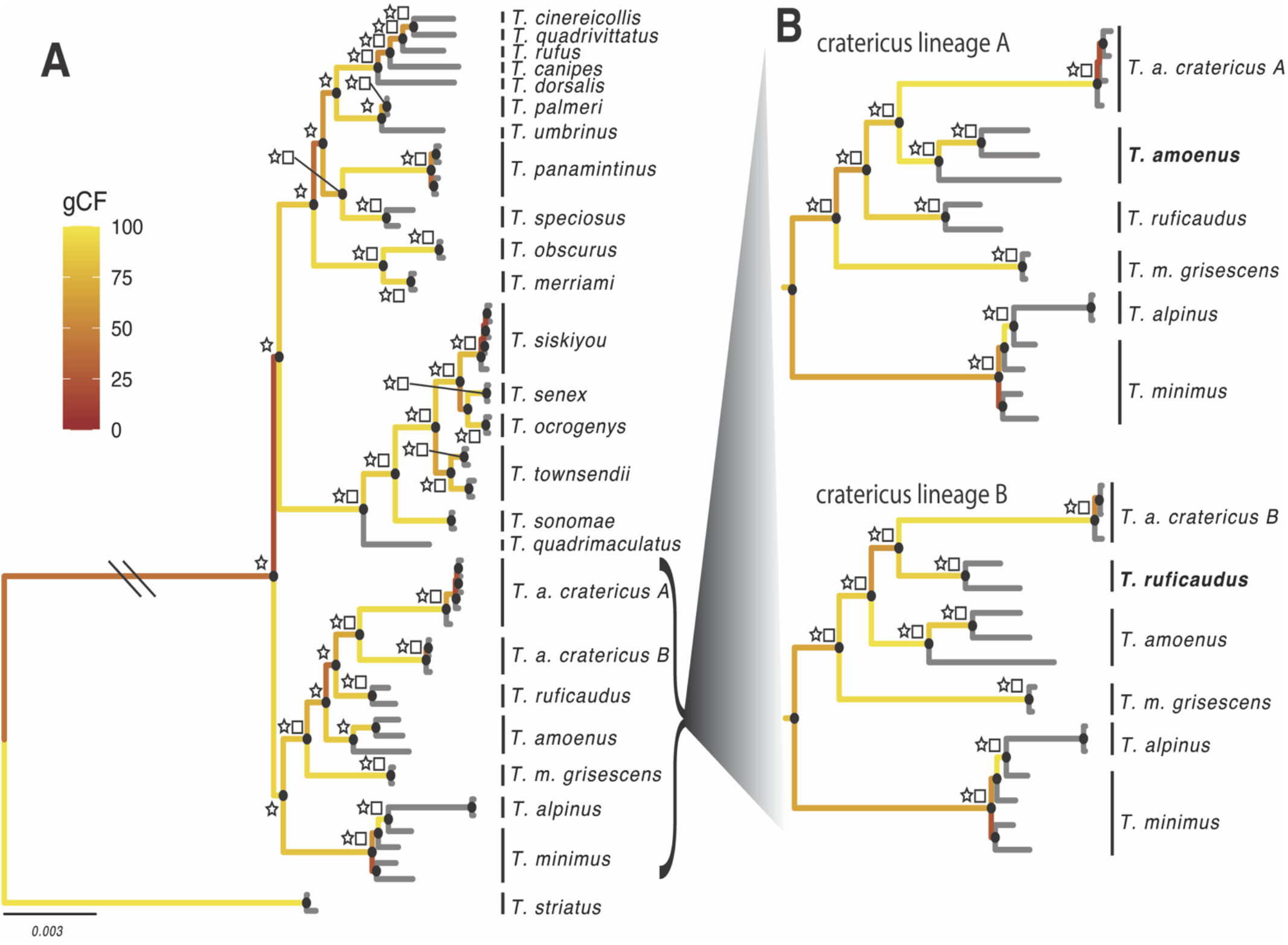
Genome-wide phylogeny of *Tamias*. **A)** Concatenated ML tree estimated with IQ-Tree; Node labels represent: ASTRAL posterior probability equal to one (star), bootstrap greater than 90 (square), and bootstrap proportions greater than 90 (grey circles). Branches are colored according to gene concordance factor (gCF). **B)** ML reconstruction of the *amoenus-ruficaudus-cratericus* clade with the exclusion of *cratericus* lineage and *cratericus* lineage respectively.

Given the deep split within the *cratericus* clade and low support for the branching structure, we hypothesized that there might be unequal ancestry between the two *cratericus* lineages, *T. amoenus,* and *T. ruficaudus*. To explore this we re-analyzed these data while excluding either *cratericus* lineage A or lineage B. When *cratericus* lineage B was excluded, we consistently recovered *cratericus* as the sister lineage to *T. amoenus* with strong support. In contrast, analyses excluding *cratericus* lineage A recovered *cratericus* as sister to *T. ruficaudus* with strong support (Figure 2B).

### POPULATION GENOMICS OF *CRATERICUS*

To elucidate the evolutionary history and geographic extent of the cryptic *cratericus* lineages, we sequenced an additional 64 chipmunks from an additional 12 localities (seven *cratericus/T. amoenus* localities; five *T. minimus* localities; Figure 3) ranging from relatively low desert sage scrub to higher elevation temperate coniferous forests throughout south central Idaho. Initial qualitative assessment of gross bacular morphologies from these samples suggested that *cratericus* may be more widespread in central Idaho, ranging north from the temperate coniferous forest/sage scrub transitional zone of the Snake River Plain to the temperate/subalpine zones of the Salmon River. Following the procedures above, we generated an alignment of 7,813,766 bp for autosomal loci and 175,884 bp for X-linked loci with >80% of individuals genotyped at a minimum coverage of 5×.

**Figure 3.**
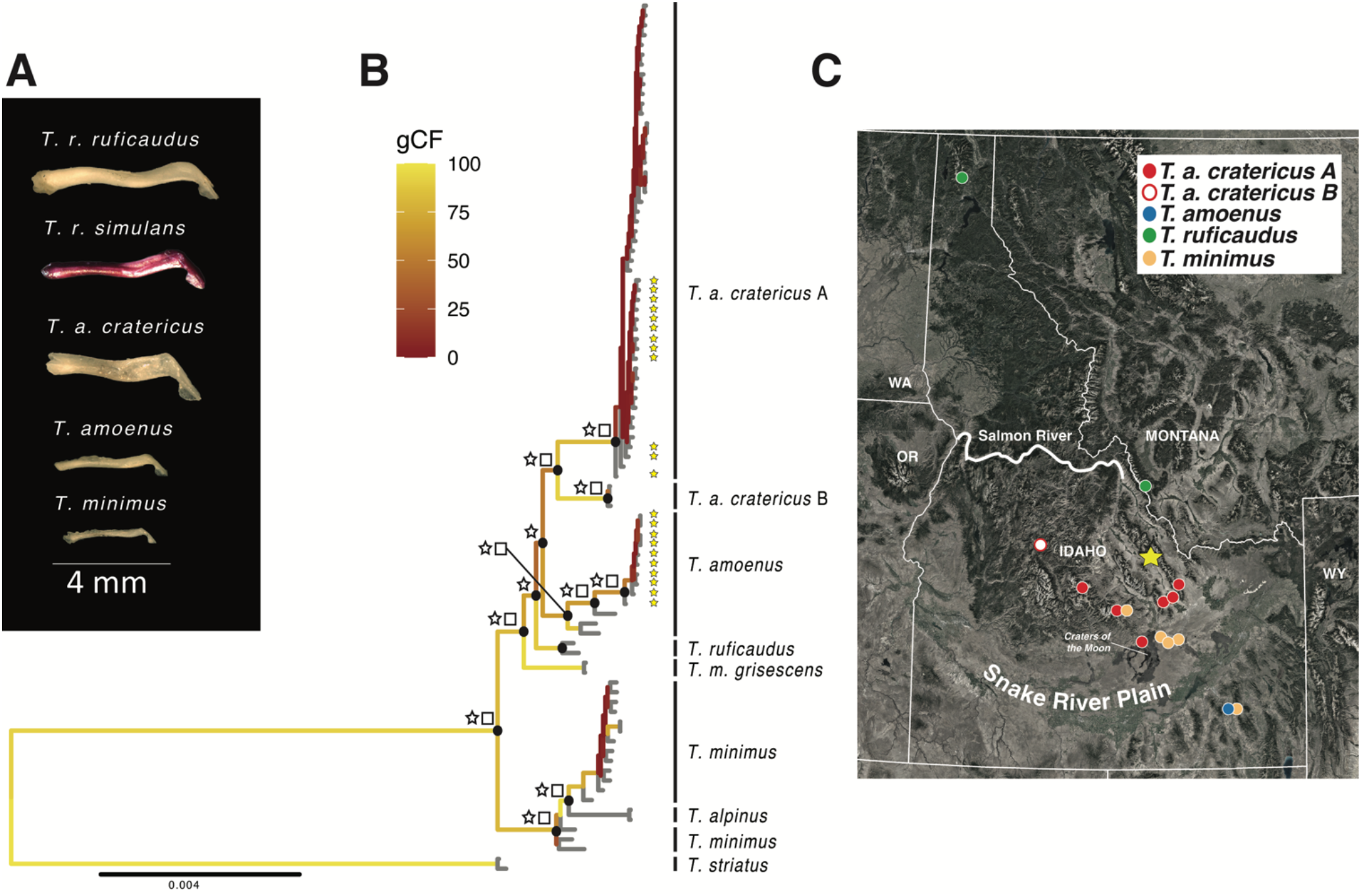
ML concatenated phylogeny for *cratericus-amoenus-ruficaudus*. **A)** Representative bacular morphologies for the five taxa. **B)** Concatenated ML tree estimated with IQ-Tree for the full population level sampling of *cratericus*; Node labels represent: ASTRAL posterior probability equal to one (star), SVDquartets bootstrap greater than 90 (square), and bootstrap proportions greater than 90 (black circles). Branches are colored according to gene concordance factor (gCF). Yellow stars at the tips indicate *cratericus* and *T. amoenus* samples from the sympatric Iron Creek locality (yellow star on map). **C)** Sampling localities for population sampling. Note that our sampling also includes non-Idaho populations for *T. minimus, T. amoenus*, and *grisescens* that are not shown.

The ML tree from the concatenated set of autosomal and X-linked loci were largely congruent with the genus-wide analysis. *T. minimus* (minus *grisescens*) was monophyletic, with *T. m. grisescens* as a distinct lineage that was more closely related to the *T. amoenus*-*cratericus*-*T. ruficaudus* complex than to other *T. minimus*. Our expanded sampling substantially increased the diversity detected within *cratericus* lineage A, which was found at 6 additional localities to the near exclusion of *T. amoenus*. We again recovered a deep split within *cratericus*, with lineage B still represented by only a single locality (Figure 3; Supplemental Figure S7).

However, the most notable difference between this expanded subset and our genus-wide analysis was that *cratericus* now appeared sister to other *T. amoenus* (Figure 3B; Supplemental Figures S10-13). One potential source of phylogenetic discordance between the datasets may be driven by samples from one locality, known as Iron Creek (IC; yellow star; Figure 3C). At this locality, both *cratericus* and standard *T. amoenus* bacular types were found to co-occur without other obvious phenotypic or habitat differences. This was the only site where both forms were found to be sympatric. Collectively, these results suggest that *cratericus* is distributed across the coniferous forest of south central Idaho, north of the Snake River Plain to the near exclusion of *T. amoenus*, save for one sympatric locality. At this site, we observed the maintenance of genetically defined species boundaries despite some evidence for admixture between *cratericus* and *T. amoenus* (see below).

Next, we evaluated if the change in the relationship between *T. ruficaudus*-*cratericus*-*T. amoenus* was influenced by possible gene flow at the sympatric IC locality. Consistent with this, *cratericus* was again sister to *T. ruficaudus* in a concatenated analysis when excluding all IC samples. However, the species trees estimated from ASTRAL (158 genes trees estimated from 50 kb genomic intervals) and SVDquartets (17,594 unlinked SNPs) for the expanded subset resulted in *cratericus* being sister to *T. amoenus*, regardless of whether IC individuals were included (Supplemental Figures S10-13). Branch support increased when IC samples were excluded but we still found a large degree of incongruence based on gCF and sCF. Finally, the results of our BPP analyses were largely consistent across the 5 replicate runs using different starting species trees, with 4 of the 5 runs converging on the same maximum *a posteriori* probability (MAP) tree, with a posterior probability of ~100%. BPP recovered *cratericus* as being sister to *T. ruficaudus* (Supplemental Figure S9).

ML reconstruction of mitochondrial genomes from the expanded subset also showed evidence for mtDNA introgression between *cratericus* and *T. amoenus* (Supplemental Figure S8). As with the genus-wide analysis, *cratericus* and *T. amoenus* were paraphyletic, but the paraphyly in the mtDNA tree was not solely driven by discordance within the IC locality. One *cratericus* mtDNA clade was composed of individuals from the majority of *cratericus* lineages A and B but excluding IC, and was sister to a monophyletic *T. minimus* clade. The second major group was composed of *T. amoenus*, including IC *T. amoenus* and *cratericus* from IC as well as a single individual from lineage B (CRCM06-160 as in the genus-wide analysis; Supplemental Figure S8).

Principal components of genetic variance (PCA) further supported a history of hybridization and mixed ancestry between *T. amoenus-cratericus*-*T. ruficaudus*. PC1 largely partitioned the most divergent lineage, *T. minimus,* from a cluster consisting of *cratericus*-*T. amoenus-T. ruficaudus*. PC2 split *T. amoenus* and *cratericus*, with *T. ruficaudus* falling intermediate. Cross-validation with ADMIXTURE suggested five population clusters that were largely concordant with the PCA (Figure 4B; Supplemental Figures S14-15), although we note that this analysis is not well-suited to partition genetic clusters at this scale of interspecific divergence (Lawson et al. 2018). Interestingly, both methods showed a tendency to cluster *cratericus* lineage B with *T. ruficaudus* and suggested a closer relationship between *cratericus* and *T. ruficaudus* than to *T. amoenus.* However, we also detected a moderate proportion of shared ancestry between sympatric (IC) *cratericus* and *T. amoenus,* likely driven by gene flow in sympatry.

**Figure 4.**
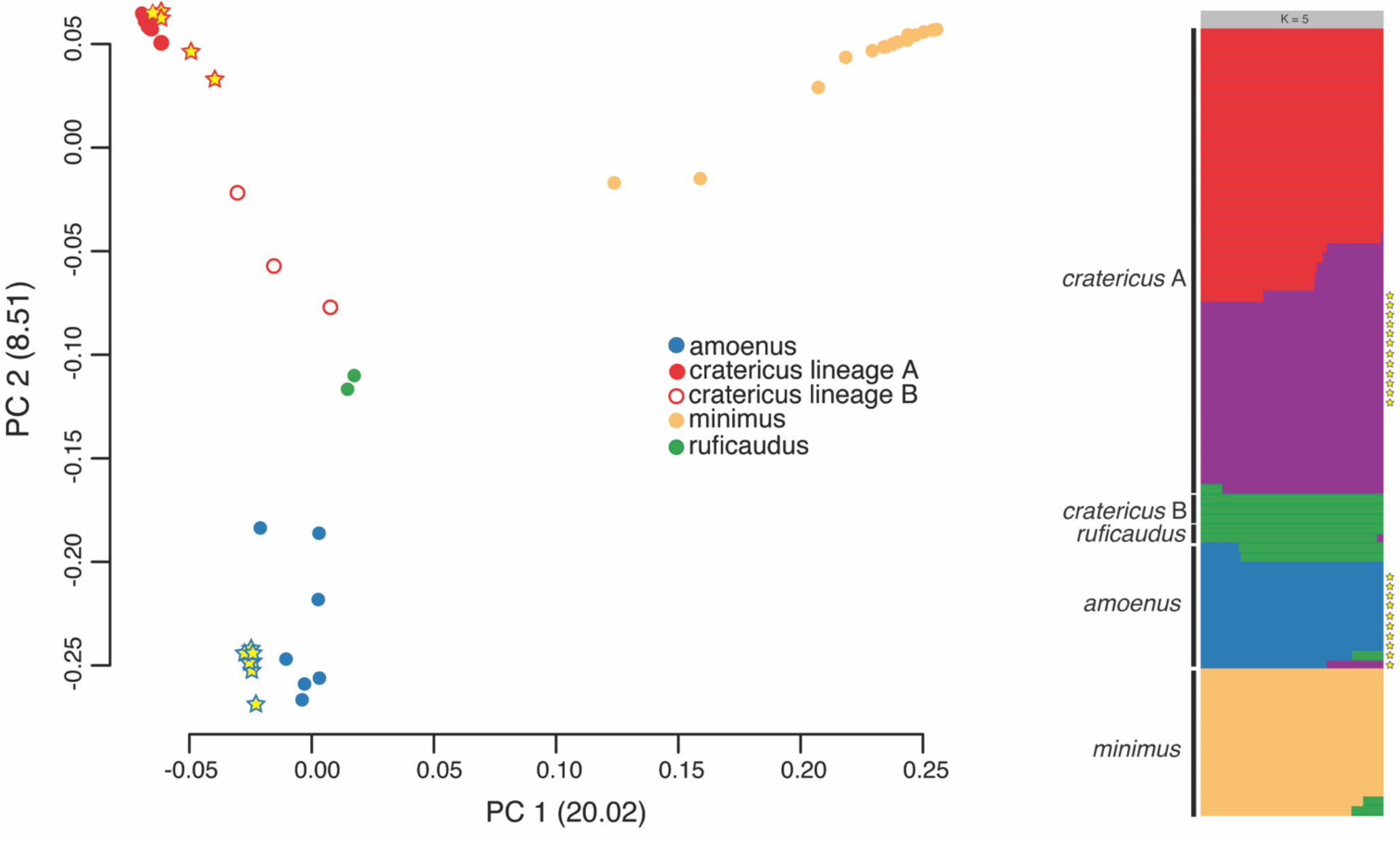
Population structure between *cratericus-amoenus-ruficaudus* sampled. **A)** PCA results from SNPRelate. **B)** Admixture output for highest marginal-likelihood run at k=5. Stars indicate samples from the sympatric locality Iron Creek.

### PATTERNS OF INTROGRESSION AND NETWORK ANALYSES

We tested for further evidence of introgression using multiple analyses based on the multispecies network coalescent (PhyloNet), quartet-based analyses under the invariants framework (HyDe) and estimates of admixture proportions based on the D-statistic (Green et al. 2010; Durand et al. 2011; Yu and Nakhleh 2015; Blischak et al. 2018). Collectively, these analyses supported varying levels of recurrent gene flow between both *cratericus*-*T. ruficaudus*, and *cratericus*-*T. amoenus*. PhyloNet supported a model of more ancient introgression, which has likely affected the inferred split between *T. amoenus-cratericus-T. ruficaudus*. The best supported network analysis (5 reticulations) suggested that *T. ruficaudus* and *cratericus* were sister lineages, with reticulation between ancestral populations of *T. amoenus* and the *cratericus* lineage. Other less well-supported reconstructions placed *T. amoenus* as more closely related to *cratericus* with a complex pattern of recurrent gene flow between *cratericus* lineage A and B with both *T. ruficaudus* and *T. amoenus* (Supplemental Figure S16). Quartet-based analyses of introgression resulted in three significant triplet comparisons supporting hybridization (Supplemental Table S4). First, *cratericus* lineage A showed shared ancestry between *T. amoenus* and *cratericus* lineage B, consistent with a close relationship between *cratericus* lineages A and B and recent hybridization between both lineages and *T. amoenus.* We also detected a pattern of shared ancestry between both *cratericus* lineages (A and B) with *T. ruficaudus* and *T. amoenus.* However, the proportion of shared ancestry was asymmetrical with respect to the amount of introgression inferred from *T. amoenus* into either *cratericus* lineage (*T. amoenus*-*cratericus* lineage A γ =0.344; *Τ. amoenus*-*cratericus* lineage B γ = 0.541). Under the ABBA-BABA framework, we evaluated phylogenetic patterns of shared SNVs between both *T. amoenus*-*cratericus* and *T. ruficaudus-cratericus.* We found that most sites supported a pattern of differential introgression between *cratericus* lineage A and both *T. amoenus* and to a lesser extent *T. ruficaudus* relative to *cratericus* lineage B (Supplemental Figure S17; Supplemental Table S4-S5).

## DISCUSSION

Discovery of cryptic species is fundamental to understanding the process of speciation and cataloging global diversity, which is facing unprecedented pressures due to the emerging threats of climate change and habitat loss. We generated a *de novo* assembled reference genome for a western chipmunk species, *Tamias minimus*, and used new and published genome-wide targeted capture sequencing data and whole mitochondrial genomes to test for cryptic speciation and hybridization in western chipmunks. Our genus-wide analysis represents the most comprehensive phylogenetic assessment of the western chipmunks to date and, we suggest, reveals at least two new undescribed chipmunk species. Below we discuss the evolutionary implications of our findings, focusing on western chipmunk diversity and the complex speciation dynamics of the Crater chipmunk and other co-distributed species.

### SYSTEMATICS OF THE WESTERN CHIPMUNK RADIATION

Due to their abundance and conspicuous nature, chipmunks have been the focus of a broad range of biological studies including physiology (e.g., Levesque and Tattersall 2009), niche partitioning and competitive exclusion (e.g., Grinnell and Storer 1924; Heller 1971), behavior (e.g. Broadbrooks 1970), and responses to climate change (e.g., Moritz et al. 2008; Bi et al. 2019). Early phylogenetic studies using morphology and allozymes (Levenson et al. 1985), host/ectoparasite (Jameson 1999), chromosomal (Nadler and Block 1962) and molecular data (Piaggio and Spicer 2000; Piaggio and Spicer 2001) recognized three distinct clades within *Tamias*: *T. sibiricus*, *T. striatus*, and the western North America species (arbitrarily classified as either subgenera or genera; (e.g., Jameson 1999; Patterson and Norris 2016). Relationships among the 23 currently recognized species within the western chipmunk clade (subgenus *Neotamias*) have remained somewhat obscure due to a lack of resolution among internal nodes and apparently rampant mitochondrial introgression (Piaggio and Spicer 2000; Reid et al. 2012; Sullivan et al. 2014). Our analysis of western chipmunks represents the most comprehensive genome-wide assessment of the group to date and included 22 of the 25 recognized species of *Tamias*.

We recovered strongly supported, monophyletic species groups within the western chipmunks that were in general agreement with some other studies (e.g., Reid et al. 2012). However, one notable difference was the relationship between *T. ruficaudus-T. amoenus-T. minimus*. In contrast to Reid et al. (2012), we recovered *T. ruficaudus* as being sister to either *cratericus*, or to a clade composed of *T. amoenus-cratericus* (see discussion below) whereas Reid et al. (2012) found *T. ruficaudus* sister to the enigmatic, and similarly distinct lineage *grisescens.* In addition, Reid et al. (2012) also recovered *T. amoenus* as sister to *T. minimus*, whereas we found *T. minimus* as the sister lineage to the main *T. amoenus-T. ruficaudus-cratericus-grisescens* clade. Given a lack of resolution among internal nodes in previous studies, this difference likely reflects the power afforded by much more extensive genetic sampling.

A major conclusion of our genomic study was the resolution of two cryptic lineages of chipmunk. While some of the relationships among described species have been unclear until recently, the identity of species as the fundamental units of diversity have remained fairly stable over the last century. The vast majority of chipmunk species were described to some degree by the late 19th and early 20th Centuries by the early naturalists Allen, Merriam, and others based on phenotypic and ecological characteristics (for a review see Thorington Jr. et al. 2012). Subsequent discussions have focused on determining if subspecific variation within some species warrant species-level recognition (Patterson 1984; Patterson and Heaney 1987; Good et al. 2003) or if some current species are actually geographically isolated populations of more widespread forms (Piaggio and Spicer 2000; Piaggio and Spicer 2001; Rubidge et al. 2014). The putative *grisescens* lineage reflects these dynamics. Howell (1925) described the distinctive *T. m. grisescens* based on its pelage as a least chipmunk subspecies restricted to the Channeled Scablands of central Washington. Reid et al. (2012) then identified this taxon as a potentially cryptic lineage and found it consistently nested with *T. amoenus* based on limited nuclear and mtDNA data. Although a formal description awaits additional geographic sampling, our data clearly show that *grisescens* does not fall within the considerable genetic diversity of the least chipmunk (*T. minimus*) or other sampled species, and we propose that *grisescens* is in fact a novel, cryptic chipmunk species (i.e., *Tamias grisescens* following Howell 1925). Of note, *grisescens* is much more genetically distinct from other chipmunk species than the Alpine chipmunk (*T. alpinus*); a lineage that is paraphyletic with respect to *T. minimus* but has long been recognized as a distinct species based on phenotypic divergence, ecological differentiation, and reproductive isolation (Grinnell & Storer; Heller; Rubidge 2014; Bi et al. 2019). We also found *grisescens* nested within *T. amoenus* in our mtDNA analyses, suggesting a history of mtDNA introgression or incomplete lineage sorting (Supplemental Figure S3). While our study lacks the sampling needed to fully investigate the evolutionary history of *grisescens* (two individuals from the same locality), its range in central Washington could contribute to vulnerability due to habitat loss, wildfire, and drought due to rapid climate change.

We also show that *cratericus* is likely a distinct chipmunk species that appears to be most closely related to *T. ruficaudus*. Blossom (1937) first described *cratericus* as a duller, dark variant of the yellow-pine chipmunk that is associated with the recent volcanic lava flows of Craters of the Moon. The color morphology of *cratericus* rapidly transitions to a more brightly colored pelage morph more like standard *T. amoenus* phenotypes and, we show, is found throughout the adjacent xeric forest habitats of the region. This localized pelage presumably reflects recent adaptation for crypsis on the black lava flows, a pattern that is also observed in local populations of the Great Basin pocket mouse (*Perognathus parvus*) and pika (*Ochotona princeps)* from Craters of the Moon (Blossom, 1937). More surprisingly, *cratericus* appears to be the locally dominant form throughout central Idaho (see below).

Overall, we also found that most western chipmunk groups displayed a high degree of discordance between the mtDNA phylogeny and the nuclear genome, consistent with previous studies documenting recurrent mtDNA introgression layered across the history of this group (e.g., Demboski & Sullivan, 2003; Sullivan et al. 2014; Sarver et al. 2017). Interestingly, previous works have suggested distinct species boundaries with comparably low levels of nuclear introgression relative to more rampant mtDNA introgression (Good et al. 2015; Sarver et al. 2021; however, see Ji et al. 2021). From this perspective, the complex evolutionary history and extensive nuclear gene flow observed between *cratericus* and both *T. amoenus* and *T. ruficaudus* is noteworthy relative to other chipmunk studies to date.

### THE COMPLEX EVOLUTIONARY HISTORY OF *TAMIAS CRATERICUS*

Our data and analyses provide compelling evidence to support *cratericus* as at least one distinct species. However, the full evolutionary history of this taxon appears to be obscured by a history of rapid speciation and likely recurrent instances of hybridization with populations of *T. amoenus* and, possibly, *T. ruficaudus*. To reconstruct this complex evolutionary history, we considered how incomplete lineage sorting and gene flow structured through space and time may lead to contrasting patterns of shared genetic variation among populations. When we assessed the placement of *cratericus* in a genus-wide context, our results suggest *cratericus* is most closely related to *T. ruficaudus* albeit with low support (Figure 2). Interestingly, SVDquartets converged on a different topology than both ASTRAL and BPP. ASTRAL and BPP agreed with the concatenated analysis and found *cratericus* sister to *T. ruficaudus,* whereas SVDquartets consistently grouped *cratericus* with *T. amoenus*. Both ASTRAL and SVDquartets appeared sensitive to sampling and the branches in question received moderate to weak support across all analyses (Figure 2; Supplemental Figures S3-S6). In contrast, while we found strong support with BPP, these analyses were limited to a relatively small subset of data which could produce misleading results given the potential for differential introgression.

Low branch support could be due to either a lack of information across all loci used to resolve relationships (e.g., due to very short internal branches and/or homoplasy) or because gene trees have independent phylogenetic histories that differ from the species tree due to incomplete lineage sorting and/or hybridization. These two sources of discordance likely compromise our ability to resolve this portion of the *Tamias* phylogeny; both gene concordance factors (gCF) and site concordance factors (sCF) for the *cratericus-T. ruficaudus* split were comparably low (gCF: 37.3; sCF: 38.6; Fig 2, Supplemental Figure S1).

One major criticism of coalescent-based summary methods, such as ASTRAL, is that they may incorrectly infer a species tree because they are sensitive to stochastic variation in phylogenetic signal between loci (Mirarab et al. 2016; Morales-Briones et al. 2018). However, the discordance observed between our approaches likely reflect biological causes of discordance such as ILS and introgression. For example, gene tree estimation error often results in a skewed ratio of gCF:sCF values (Minh et al. 2020a), a pattern we do not see, even across the weakest supported branches.

While individual tree-based approaches may be limited when local genealogies are poorly resolved due to rapid divergence or limited data, our data also strongly supported the conclusion that much of the phylogenetic discordance likely reflects introgressive hybridization. We consistently detected genealogical asymmetry between *cratericus* lineage A and *cratericus* lineage B with respect to both *T. ruficaudus* and *T. amoenus*. When we assessed the relationship of either *cratericus* lineage in the absence of the other, we found strong support for two different topologies. When we excluded *cratericus* lineage B, *cratericus* lineage A was consistently sister to *T. amoenus*. Conversely, when we excluded *cratericus* lineage A, we recovered *cratericus* lineage B as the sister group to *T. ruficaudus* (Figure 2; Supplemental Figures S8-S9). Further, we inferred multiple reticulations across the *amoenus-cratericus-ruficaudus* splits, suggesting there has been both recent and ancient hybridization between these lineages during their diversification. Both contemporary and past hybridization events have resulted in extensive shared polymorphism among the three lineages (Supplemental Figures. S16-S17) and highlight how rapid diversification and introgressive hybridization can confound our ability to infer a complex evolutionary history among species.

Integrating our phylogenetic and population genomic data, we propose a working model for the evolutionary history of *cratericus* (Figure 5). There was an initial (likely geographic) split between *cratericus* and other *T. ruficaudus*, followed by intermittent hybridization between ancestral populations of *cratericus, T. ruficaudus,* and the slightly more distantly related *T. amoenus.* Determining if the apparent genetic split within *cratericus* is associated with reproductive isolation between these lineages or divergence in other morphological or ecological traits awaits further sampling. While lineage A appears to be the widespread and predominant form, the rarity of lineage B may be due to a lack of sampling in the presumed northern range of *cratericus.* We also detected gene flow between *T. amoenus* and the more widely distributed *cratericus* lineage A (Figures 3–4). While a history of recurrent hybridization likely skews overall phylogenetic relationships, the sympatric population (IC) allows us to test key predictions about the extent of reproductive isolation between these ecologically and phenotypically similar species. In this area of sympatry, individuals clearly clustered within *cratericus* lineage A or with other *T. amoenus*. Thus, species boundaries appear to be largely maintained, despite some level of gene flow between *cratericus* and *T. amoenus* (Figures 3–4; Supplemental Figure S14), consistent with a semipermeable species boundary (Harrison and Larson 2014). Our results also indicate that *cratericus* is the predominant chipmunk species associated with xeric forests of south-central Idaho north of the Snake River Plain and south of the Snake River. This previously unknown lineage appears to exist largely to the exclusion *T. amoenus.* Since the foundational work of Grinnell a century ago (Grinnell and Storer 1924), western chipmunks have been considered exemplars of ecological niche partitioning. Given the lack of obvious ecological and phenotypic differentiation between *cratericus* and *T. amoenus*, sympatric populations between these species may provide a compelling eco-evolutionary context to understand both the local maintenance of species boundaries and niche partitioning through competitive exclusion or other mechanisms.

**Figure 5.**
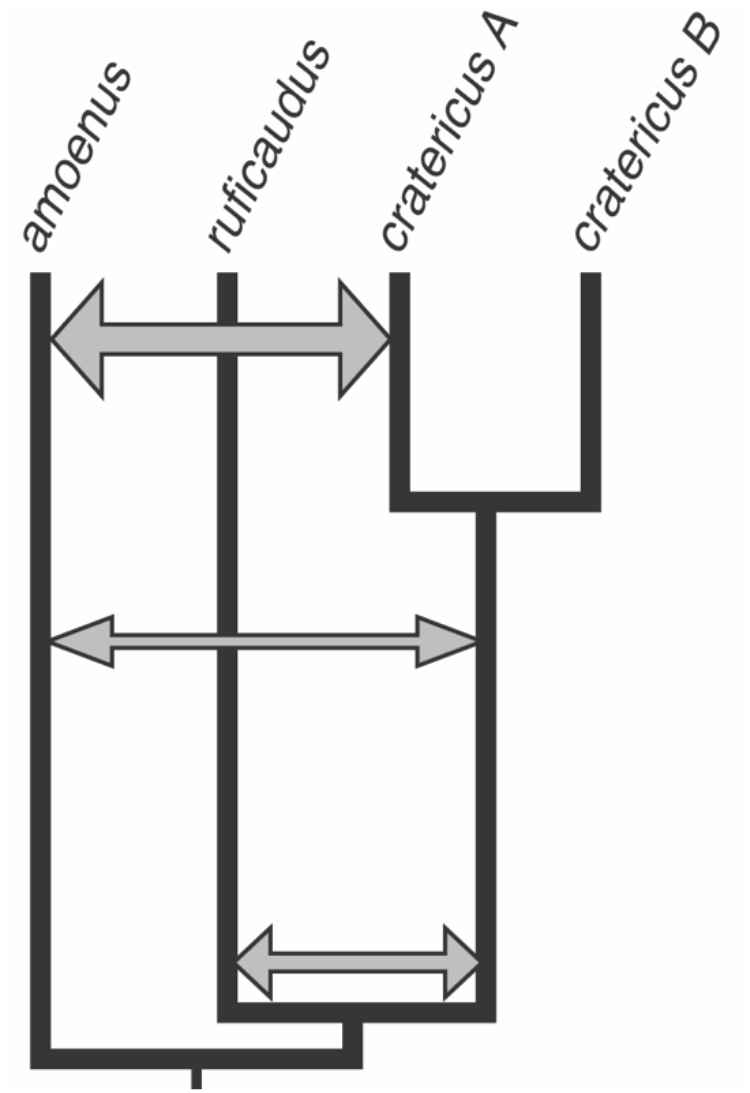
Proposed species tree and evolutionary history of *T. cratericus*. *T. cratericus* has a complex evolutionary history punctuated by layered ancient and contemporary gene flow between co-distributed species. Arrow thickness indicates inferred amount of introgression.

Finally, our current sampling is not circumscribed by clear geographic barriers to the north and west, indicating the full range of *cratericus* may be considerably larger and likely extends into the remote wilderness tracks of central Idaho. Although these portions of the range are more difficult to access, this diurnal cryptic species is also highly abundant and readily visible around the visitor center of a National Monument that hosts ~200,000 annual visitors – underscoring that cryptic biodiversity may persist even in highly conspicuous systems.

## Data Archive

Read sequence data are available for download at SRA under the BioProject accession number XXXX.

## Author contributions

NDH, JRD, JS, KCB, and JMG conceived and designed the study. JMG, JRD, and JS acquired funding. NDH, KCB, JRD, and JMG conducted fieldwork. CMC, EN, and NDH generated sequence data. NDH conducted data analyses, with guidance from BAJS and JMG. BAJS contributed analytical tools. All authors discussed the results. NDH and JMG wrote the manuscript with feedback from all authors.

## Acknowledgements

Our research would not be possible without the irreplaceable support of natural history museums. We are grateful to the collections staff of The Denver Museum of Nature & Science and Joseph A. Cook and the University of New Mexico Museum of Southwest Biology for providing tissue loans. We also thank Michael Fazekas, Roger Rodriguez, Patricia McDonald, Randle McCain, Bryan McLean, Schuyler Liphardt, and Lois Alexander for help with fieldwork. We thank David Xing, Jessi Kopperdahl, Mickael Fazekas, and Sara Keeble for assisting with molecular work. We thank members of the Good lab, the University of Montana UNVEIL network, Ke Bi, and Craig Moritz for helpful discussions. Funding support for this research was provided by a grant from the National Science Foundation (NSF) EPSCoR (OIA-1736249 to JMG), NSF (DEB-0716200 to JRD), a travel grant from the Drollinger-Dial Foundation, the Gordon and Betty Moore Foundation (GBMF2983), the Rose Community Foundation, and research funds from the University of Montana and Denver Museum of Nature & Science. This study included research conducted in the University of Montana Genomics Core, supported by a grant from the M. J. Murdock Charitable Trust (to JMG). Computational resources and support from the University of Montana’s Griz Shared Computing Cluster (GSCC), supported by grants from the Nation Science Foundation (CC-2018112 and OAC-1925267), contributed to this research. The DNA isolation, library preparation, and sequencing of the draft *Tamias minimus* genome was carried out at the DNA Technologies and Expression Analysis Core at the UC Davis Genome Center, supported by NIH Shared Instrument Grant 1S10OD010786-01.

