## Supplemental Figures for "Genomic resolution of cryptic species diversity in chipmunks"

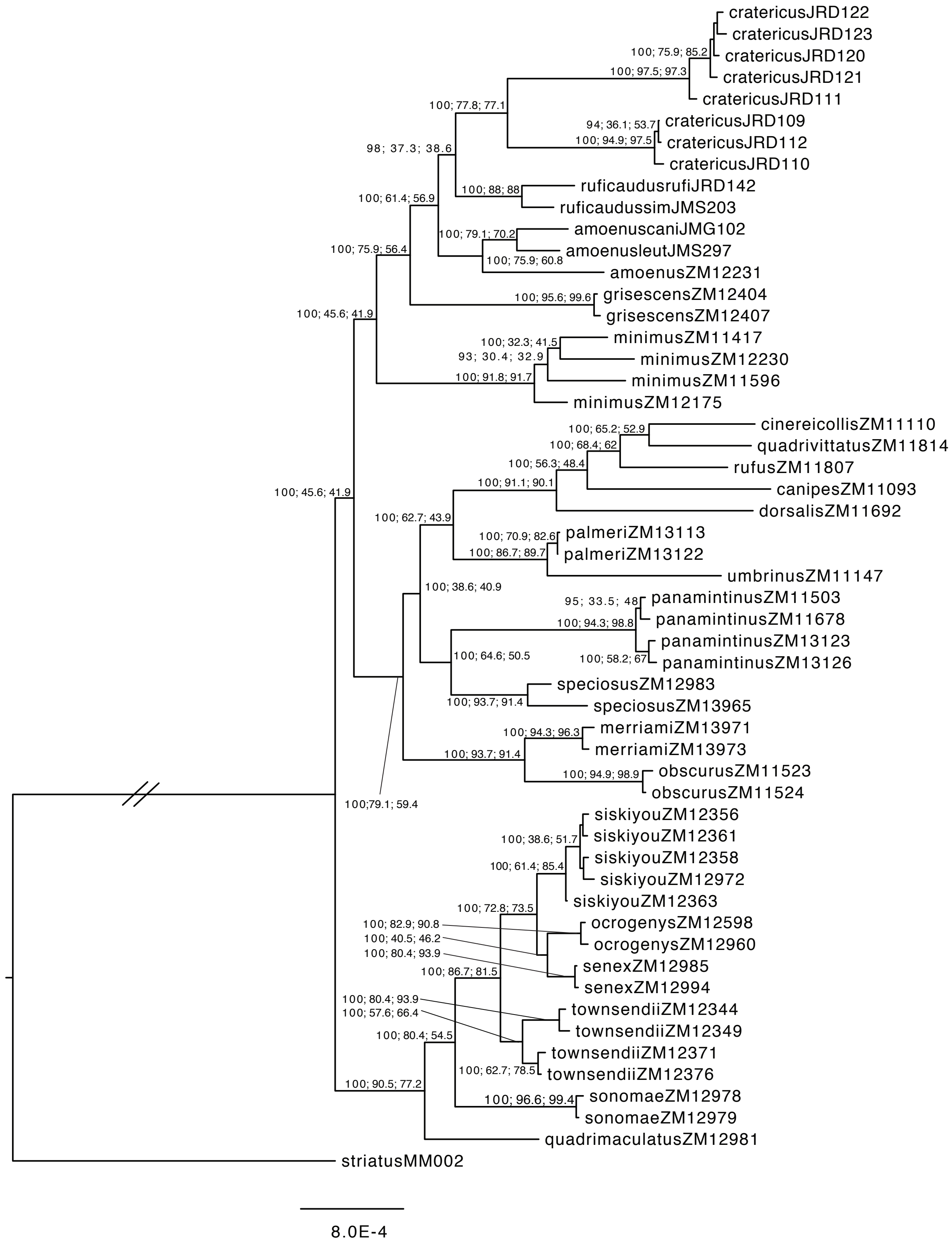


Figure S1 – Maximum likelihood phylogram of concatenated dataset (5,365,556 bp) for the total Tamias tree (54 Ind.). To accommodate the smaller (~6 Mb) capture of Sarver et al. (2021), we subset our larger capture (~9.2 Mb) to mitigate sequencing depth biases between the captures. Node labels indicate UFBoot > 80%, gCF, sCF values.

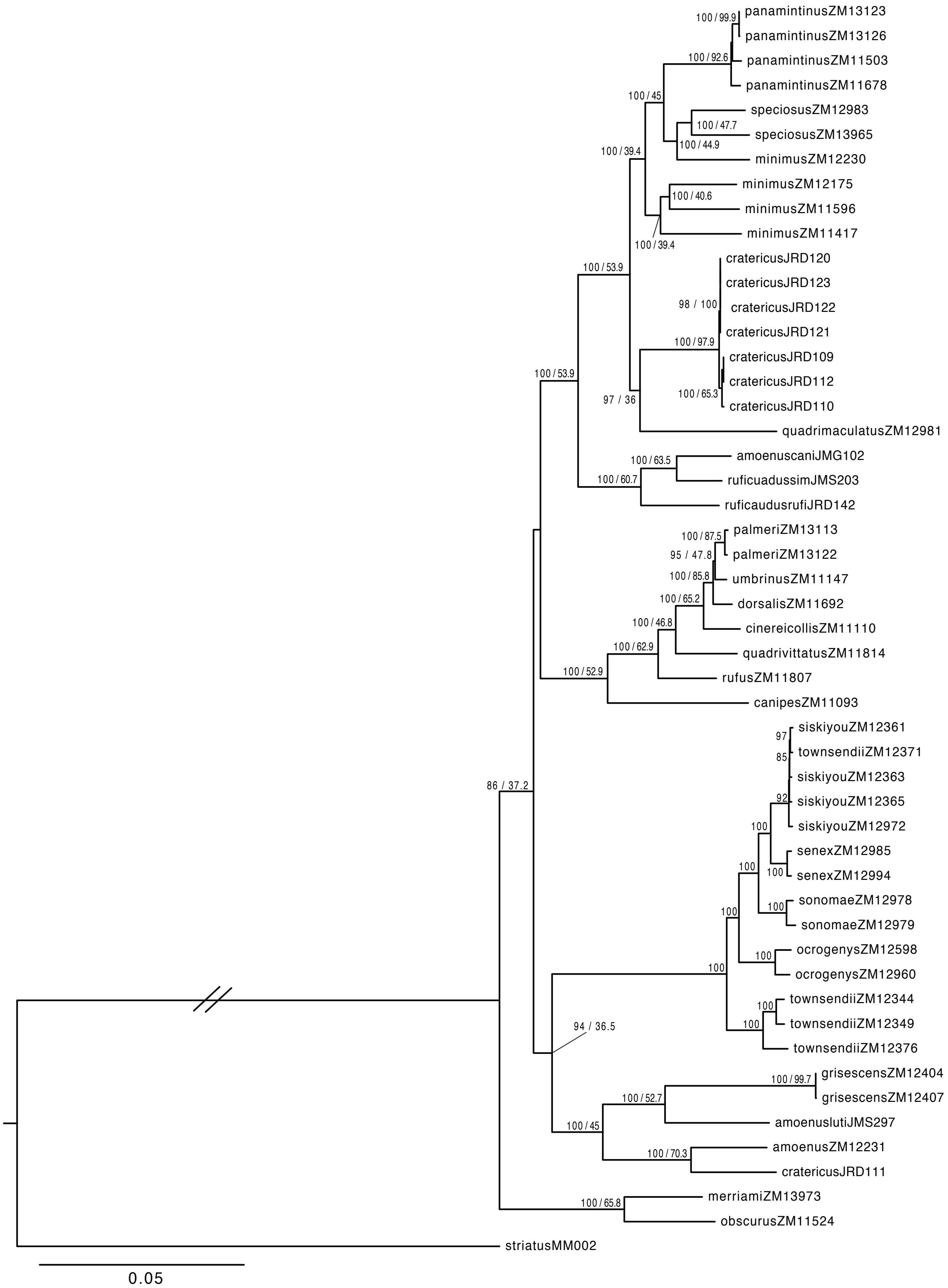


Figure S2 – Maximum likelihood phylogram of mitochondrial genomes (16,697 bp). Node labels indicate UFBoot values of > 80% and site concordance factors (sCF).

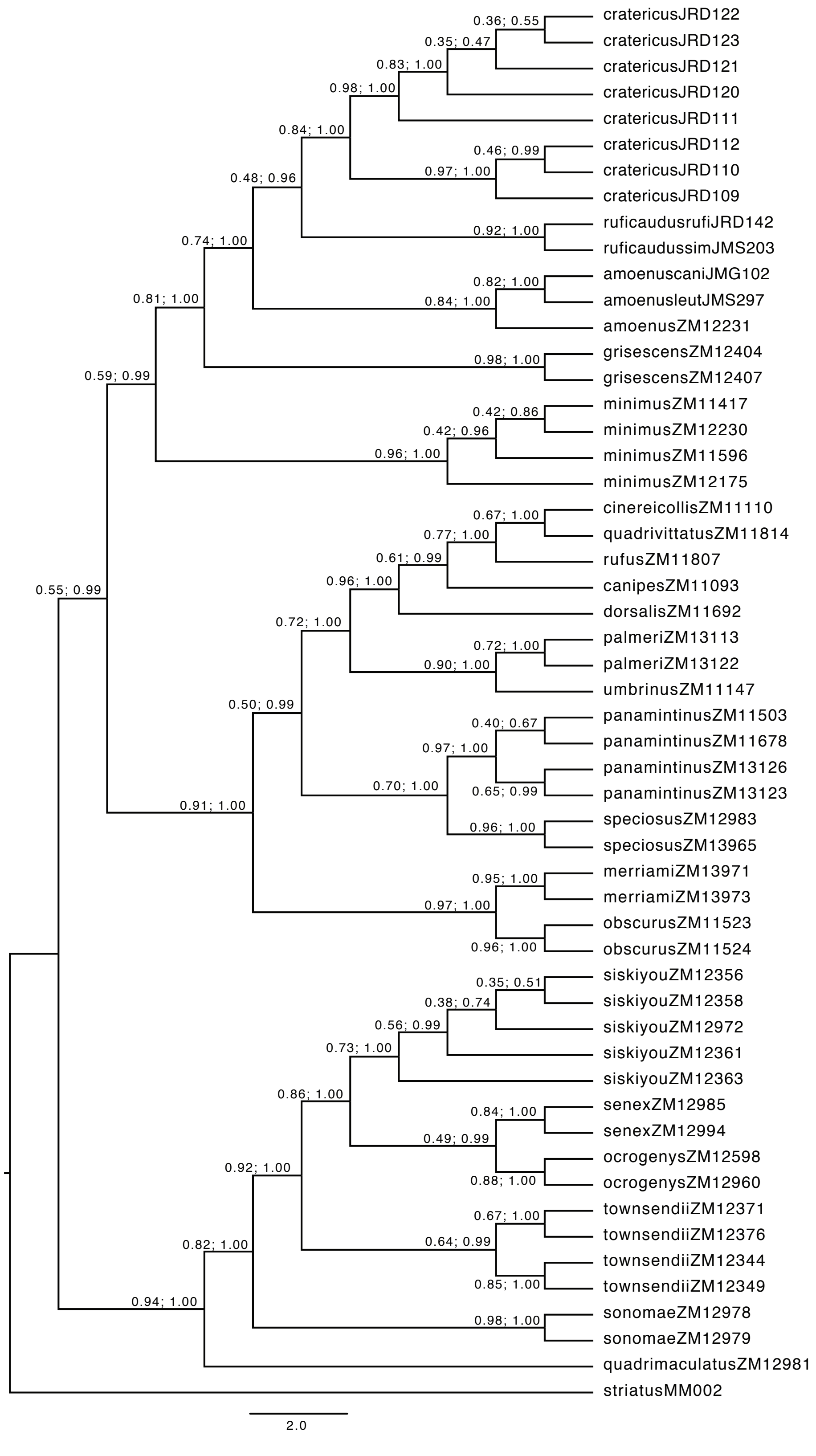


Figure S3 – ASTRAL species tree obtained from 158 gene trees without assignment of individuals to species. Nodes are annotated with quartet scores and posterior probabilities.

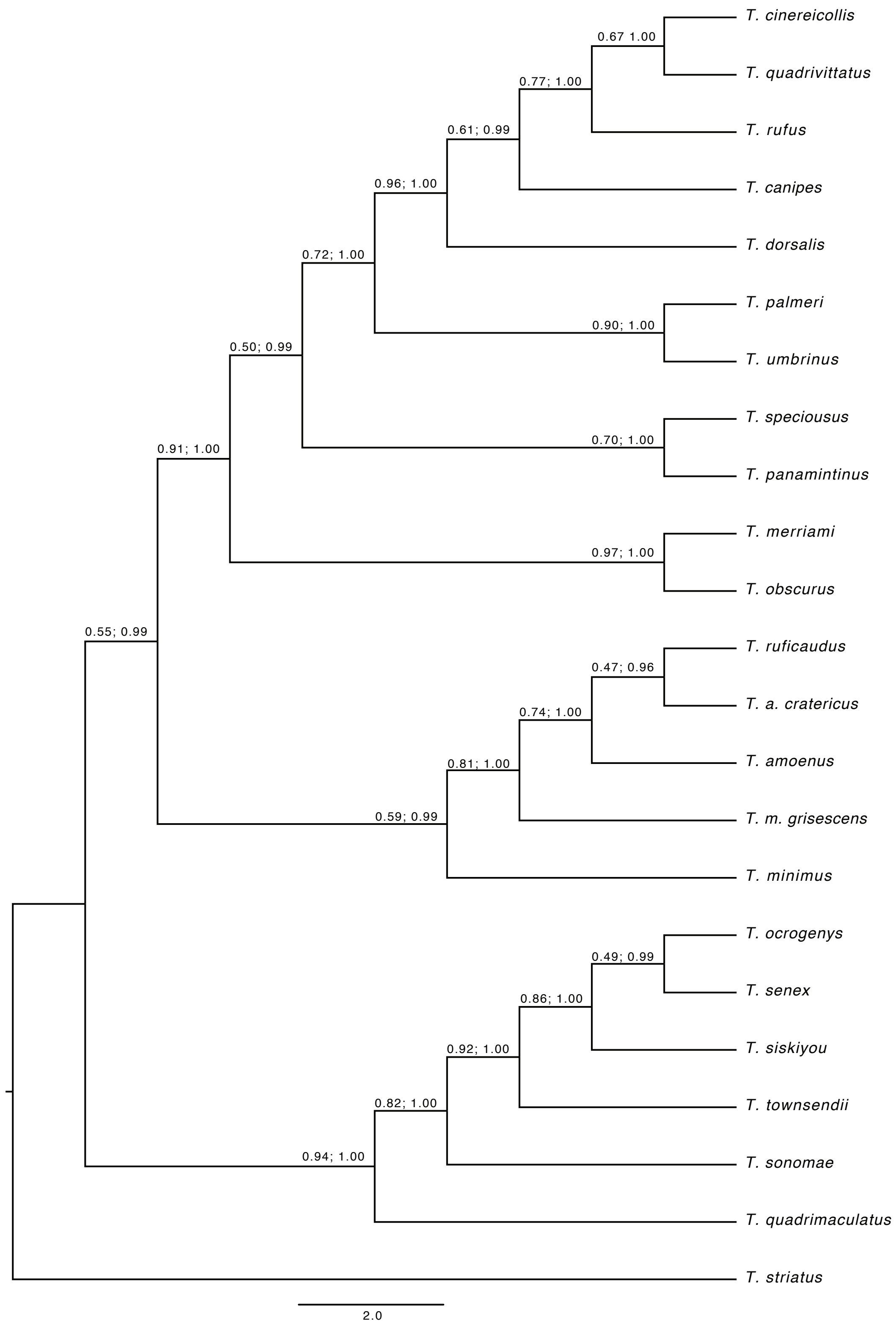


Figure S4 – ASTRAL species tree obtained from 158 gene trees with assignment of individuals to species. Nodes are annotated with quartet scores and posterior probabilities.

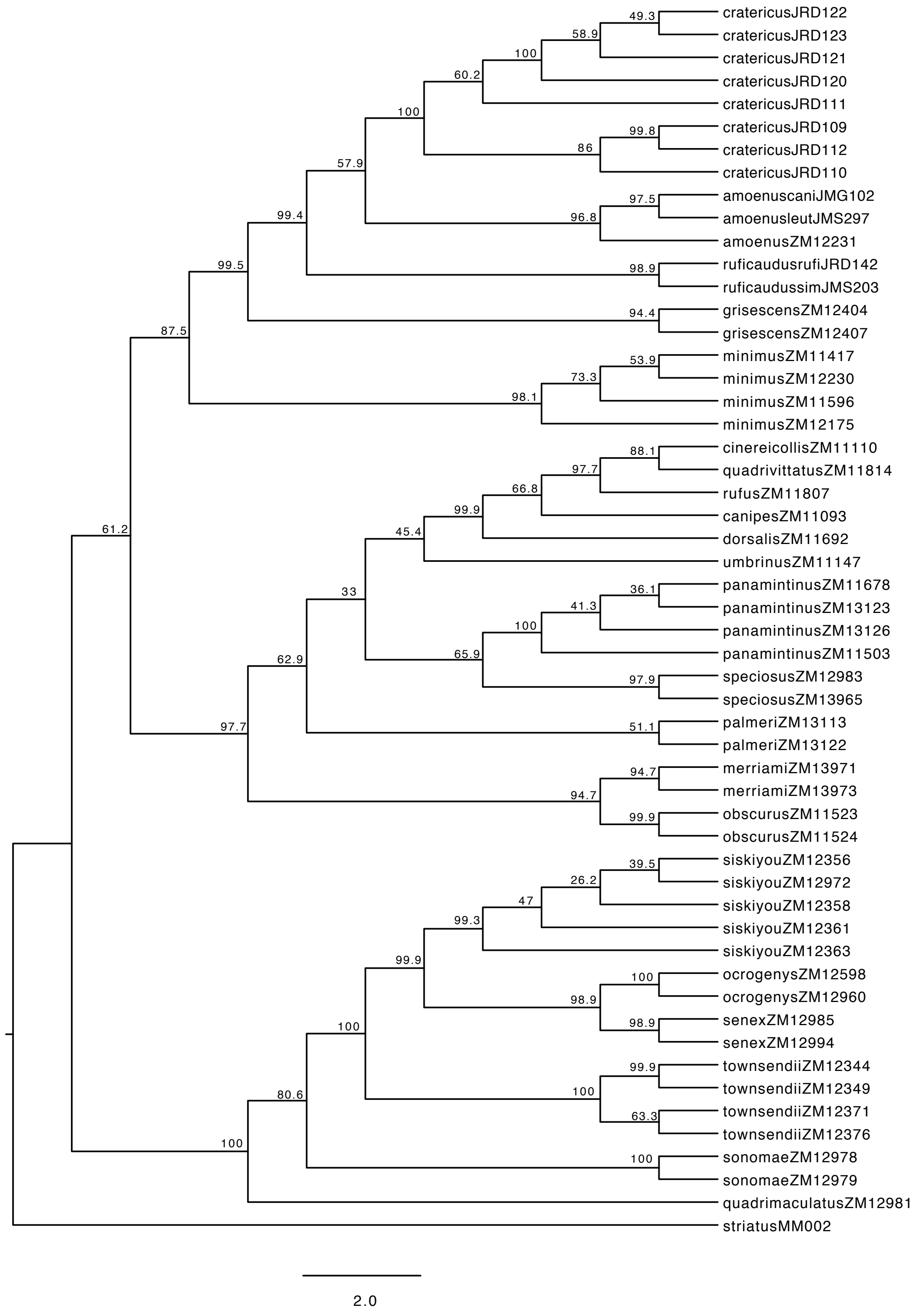


Figure S5 – *SVDquartets* species tree generated with 13,482 unlinked SNPs without assignment of individuals to a species. Branches are labeled with bootstrap proportions from 1000 bootstrap iterations.

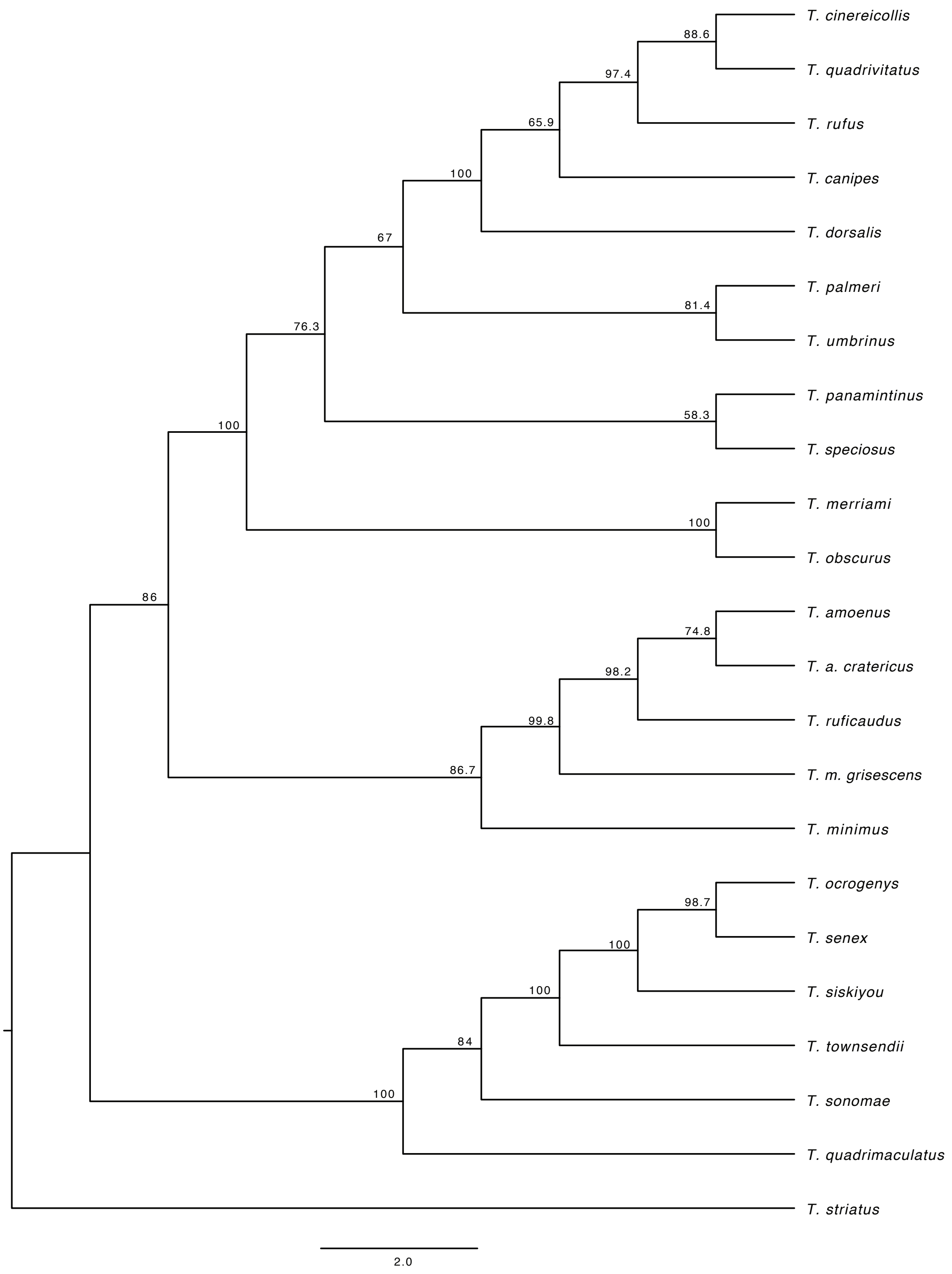


Figure S6 – *SVDquartets* species tree generated with 13,482 unlinked SNPs with assignment of individuals to species. Branches are labeled with bootstrap proportions from 1000 bootstrap iterations.

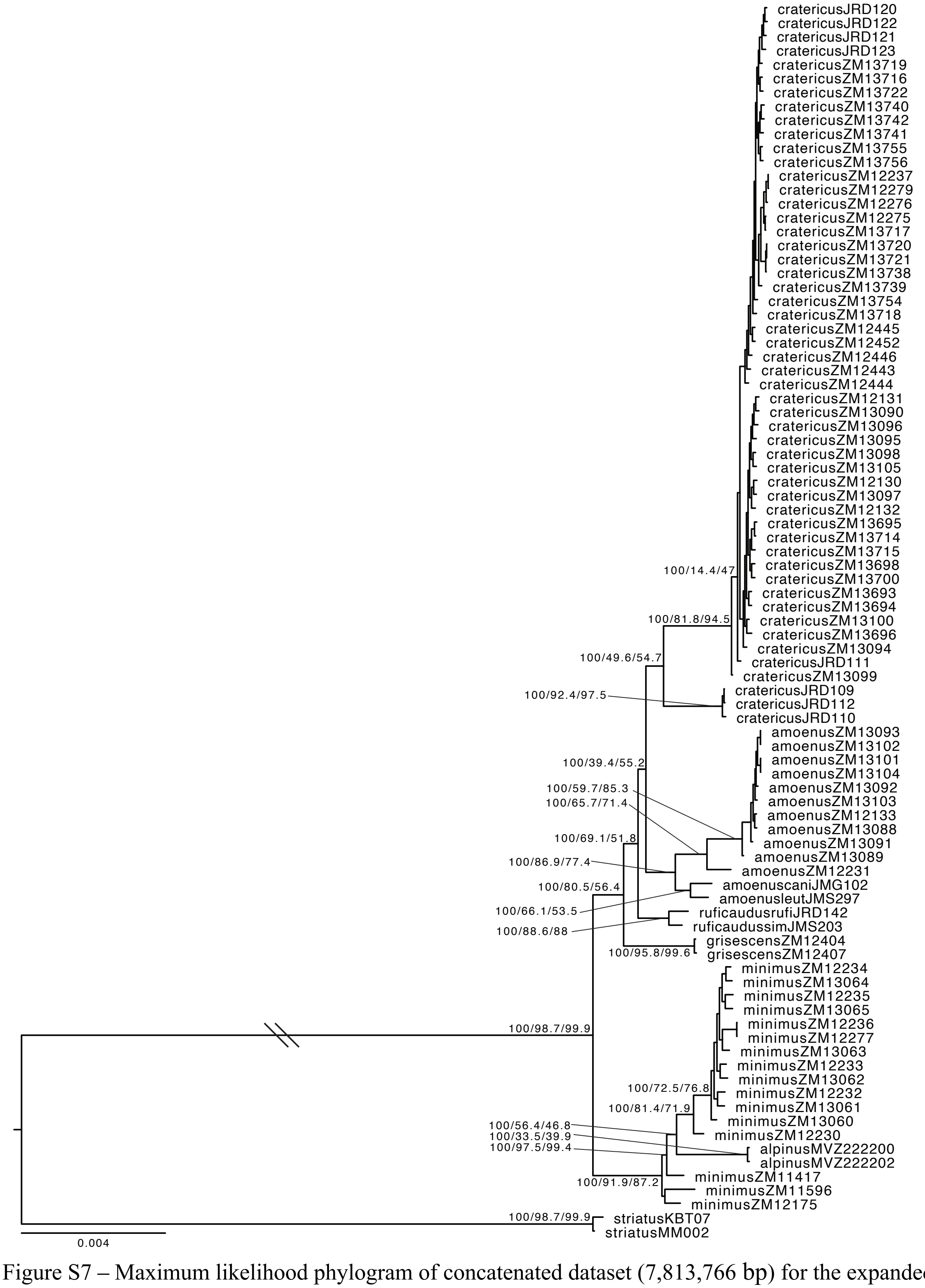


Figure S7 – Maximum likelihood phylogram of concatenated dataset (7,813,766 bp) for the expanded Tamias subset tree (84 Ind.). Branch labels represent UFBoot, gCF, and sCF values and are shown for major nodes only.

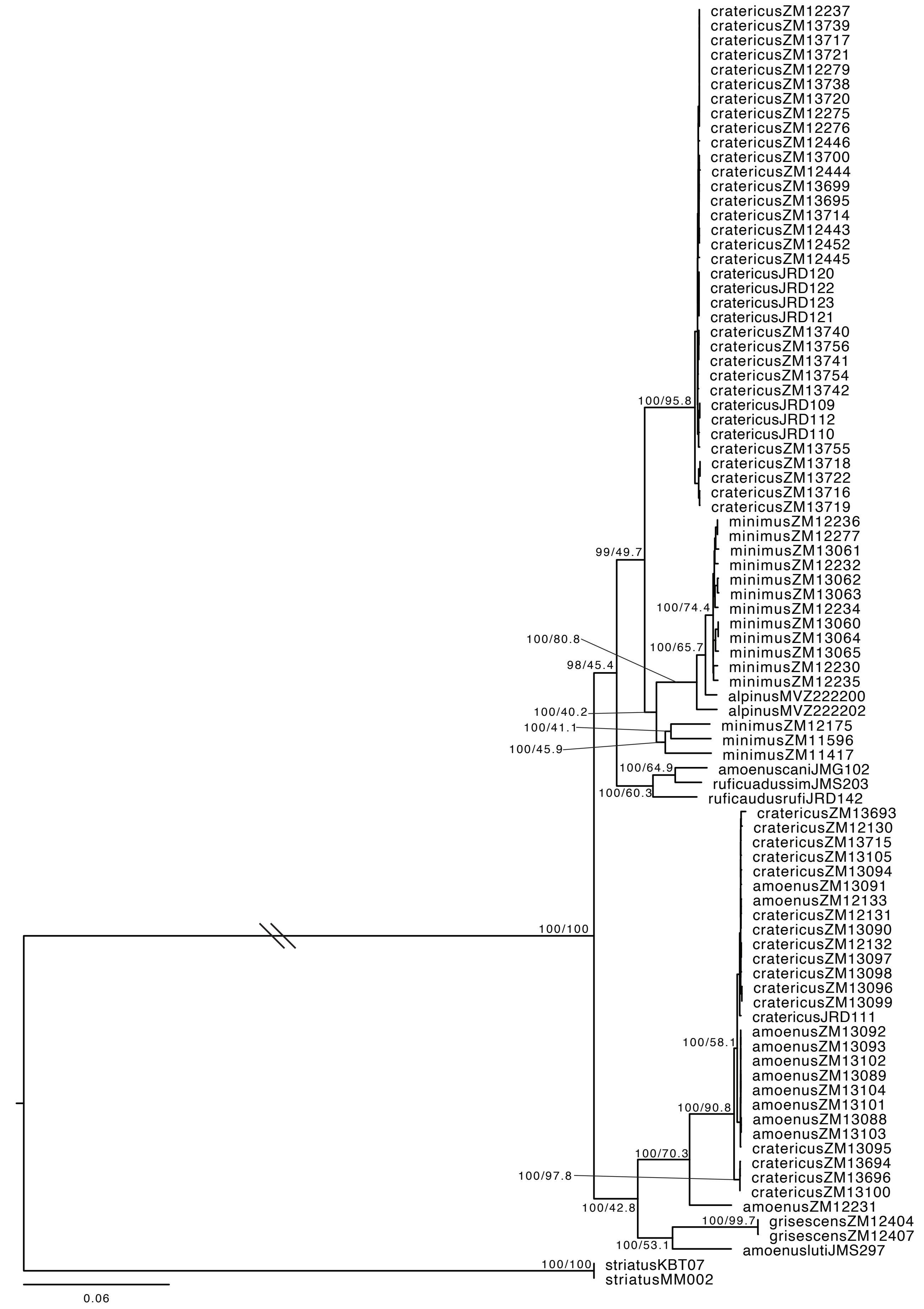


Figure S8 – Maximum likelihood phylogram of mitochondrial genomes (16,697 bp) for the expanded Tamias subset tree (84 Ind.). Node labels indicate UFBoot values of > 80% for major splits.

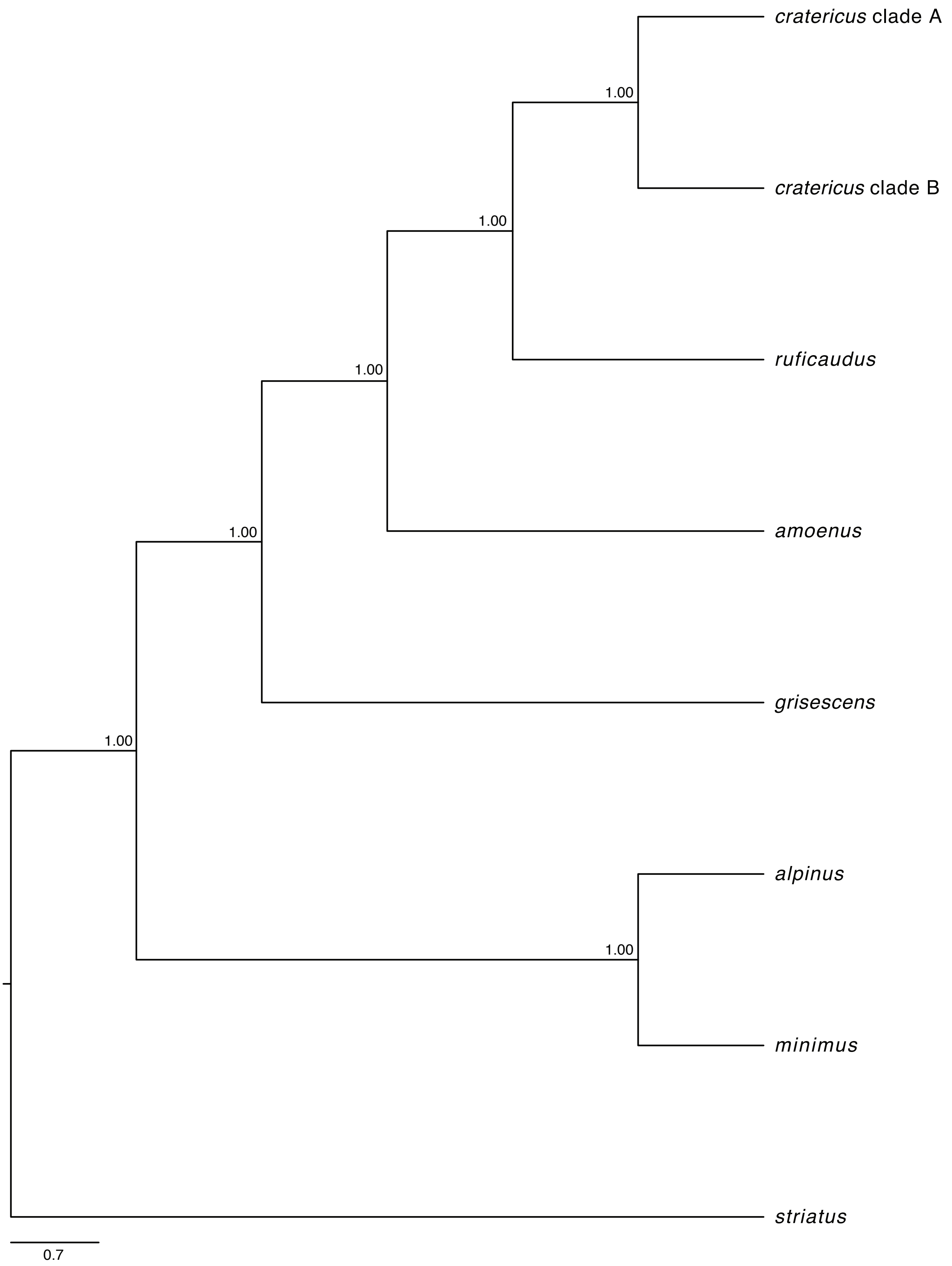


Figure S9 – BPP species tree obtained from 88 loci (~300 kbp) with assignment of individuals to species. Nodes are annotated with posterior probabilities.

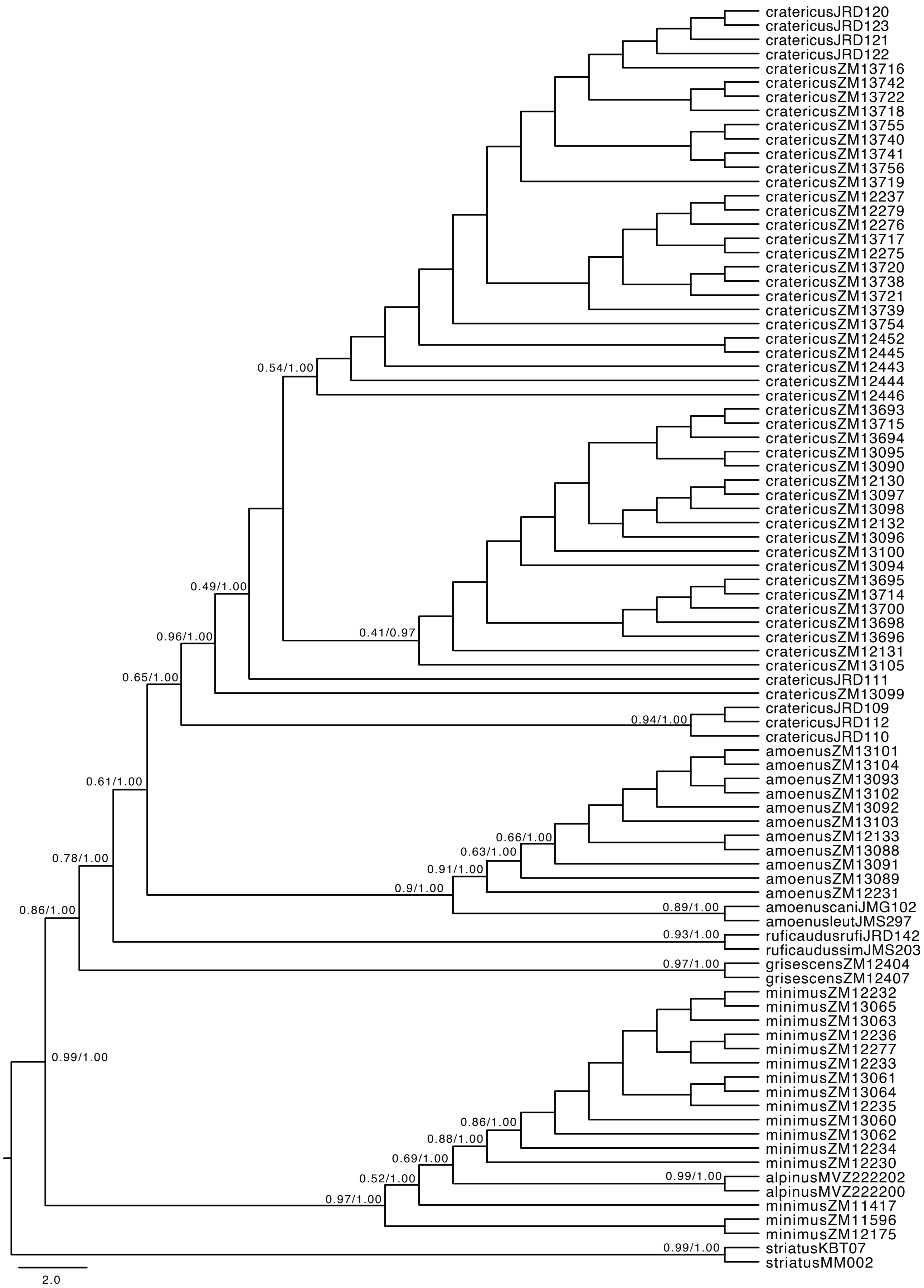


Figure S10 – ASTRAL species tree obtained from 161 gene trees without assignment of individuals to species. Nodes are annotated with quartet scores and posterior probabilities.

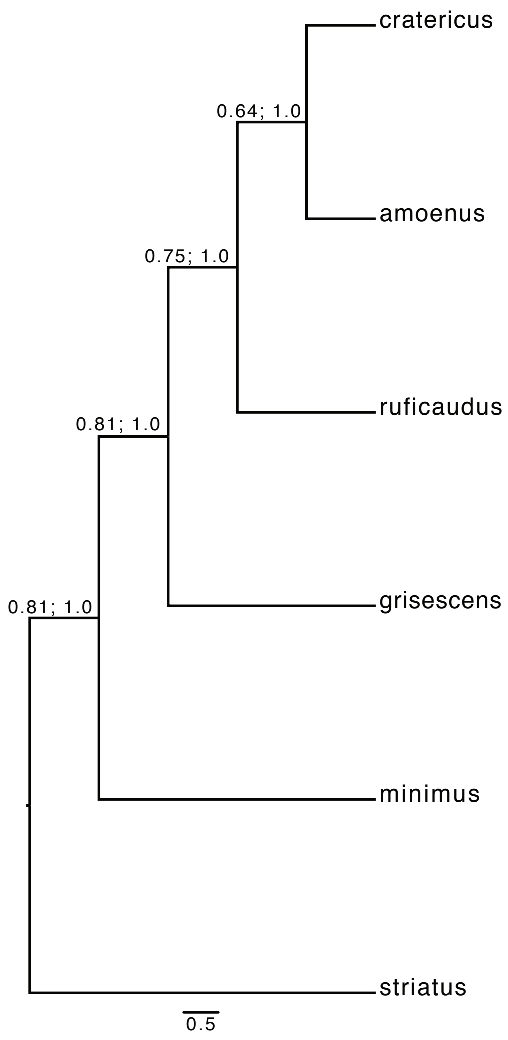


Figure S11 – ASTRAL species tree obtained from 161 gene trees with assignment of individuals to species. Nodes are annotated with quartet scores and posterior probabilities.

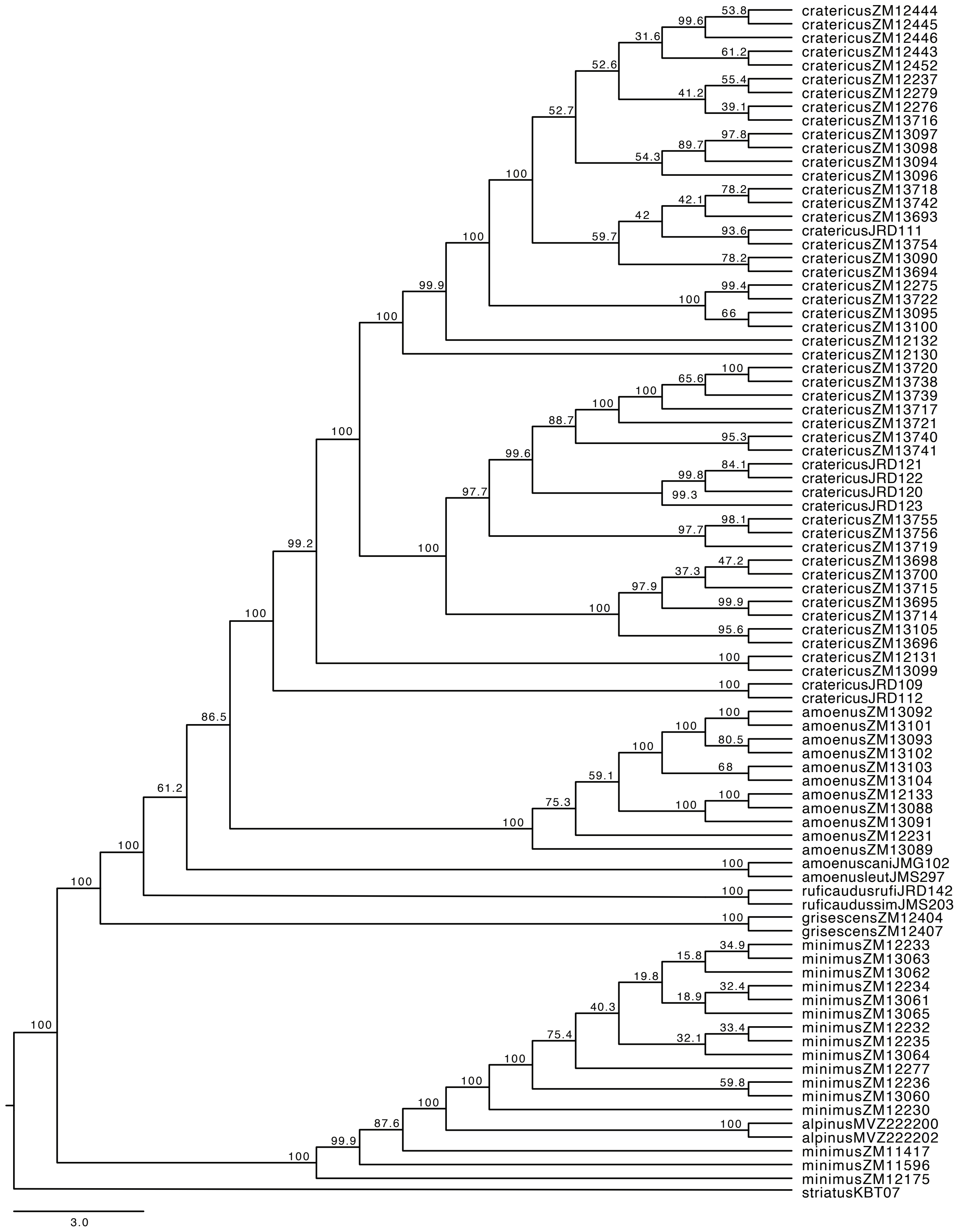


Figure S12 – *SVDquartets* species tree generated with 17,594 unlinked SNPs without assignment of individuals to a species. Branches are labeled with bootstrap proportions from 1000 bootstrap iterations.

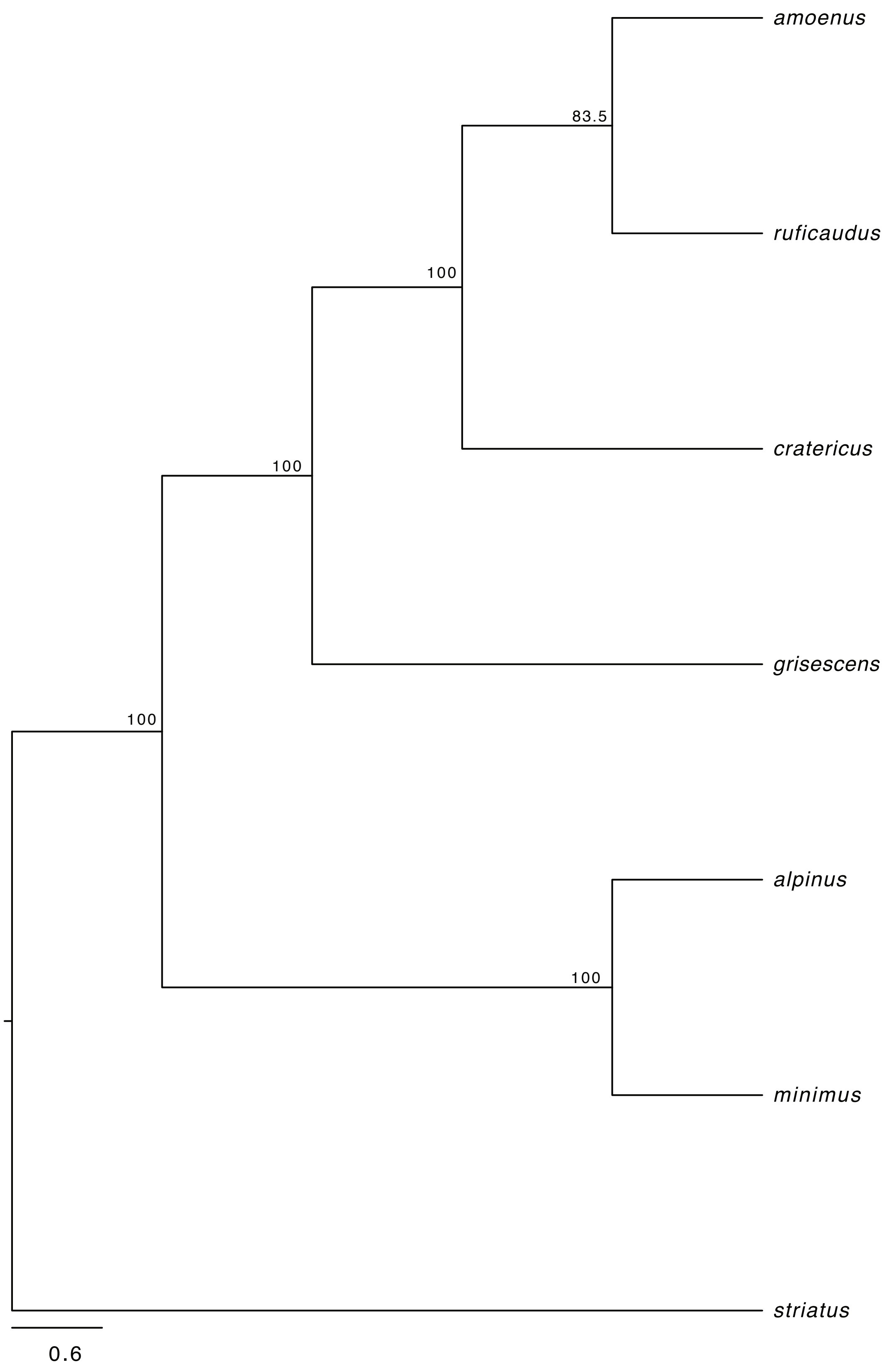


Figure S13 – *SVDquartets* species tree generated with 17,594 unlinked SNPs with assignment of individuals to species. Branches are labeled with bootstrap proportions from 1000 bootstrap iterations.

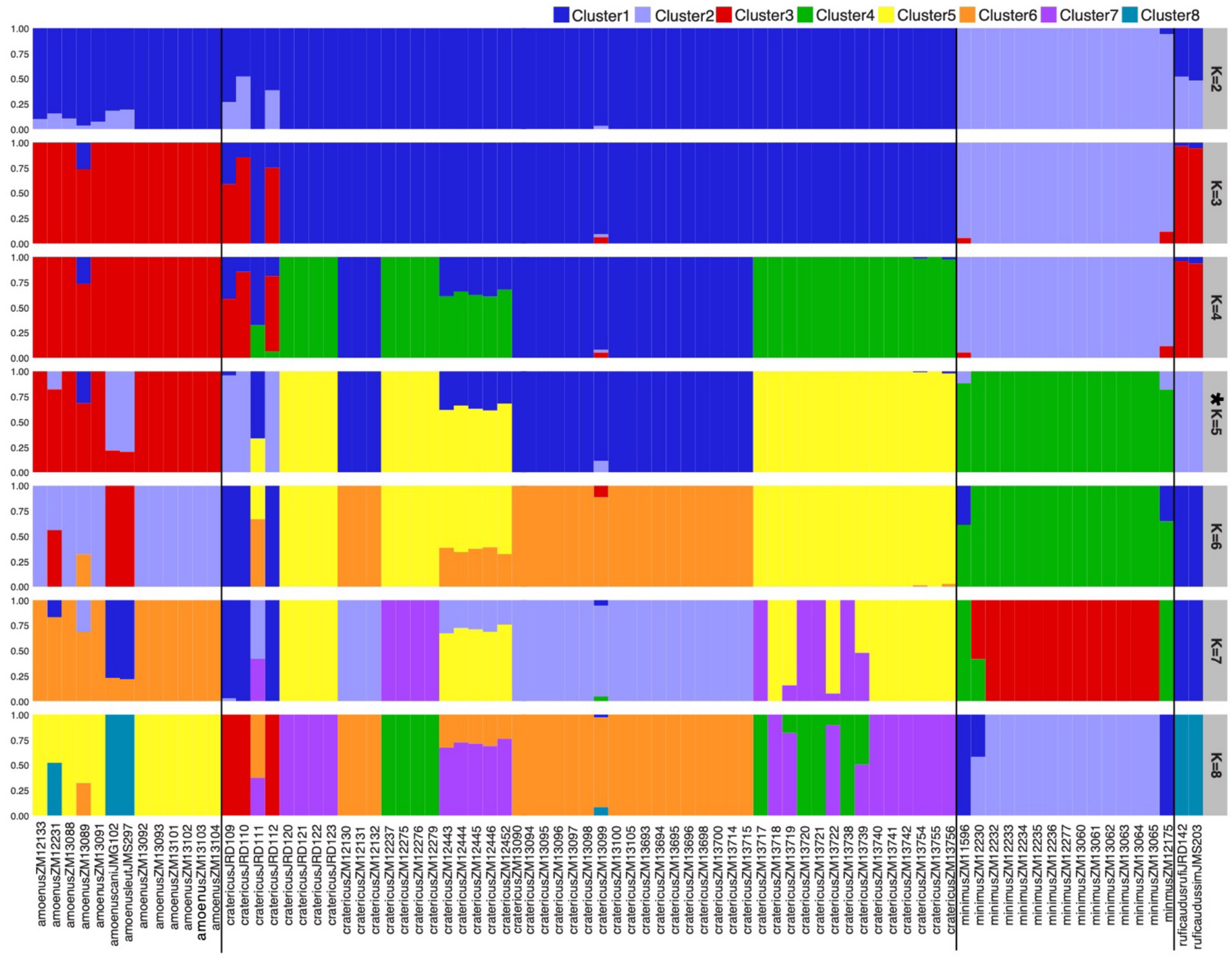


Figure S14 – Clustering of individuals in ADMIXTURE analyses with varying number of clusters for K = 2 to K = 8. Plots are organized by species designation (indicated by solid black line). K = 5 was the optimal number of K clusters based on Cross-validation.

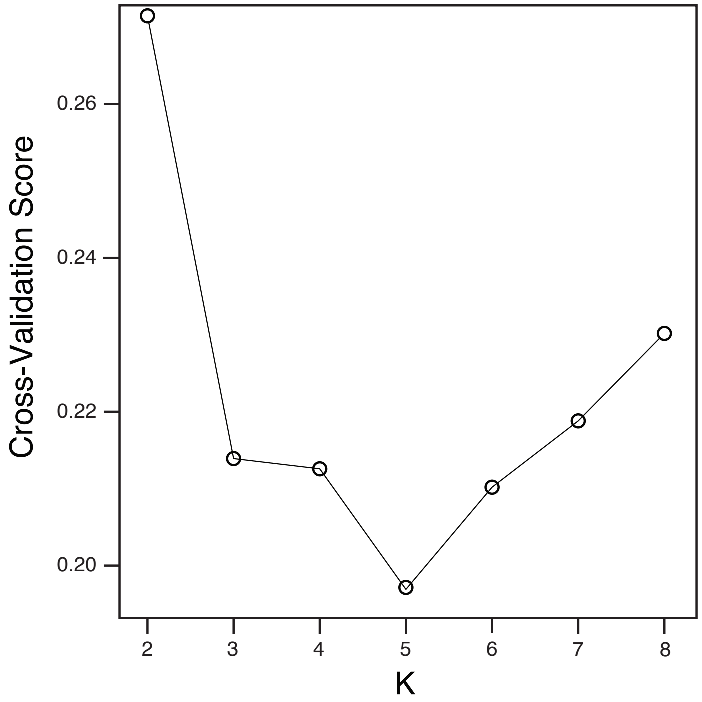

Figure S15 – Cross validation of number of populations (K) in ADMIXTURE analyses.

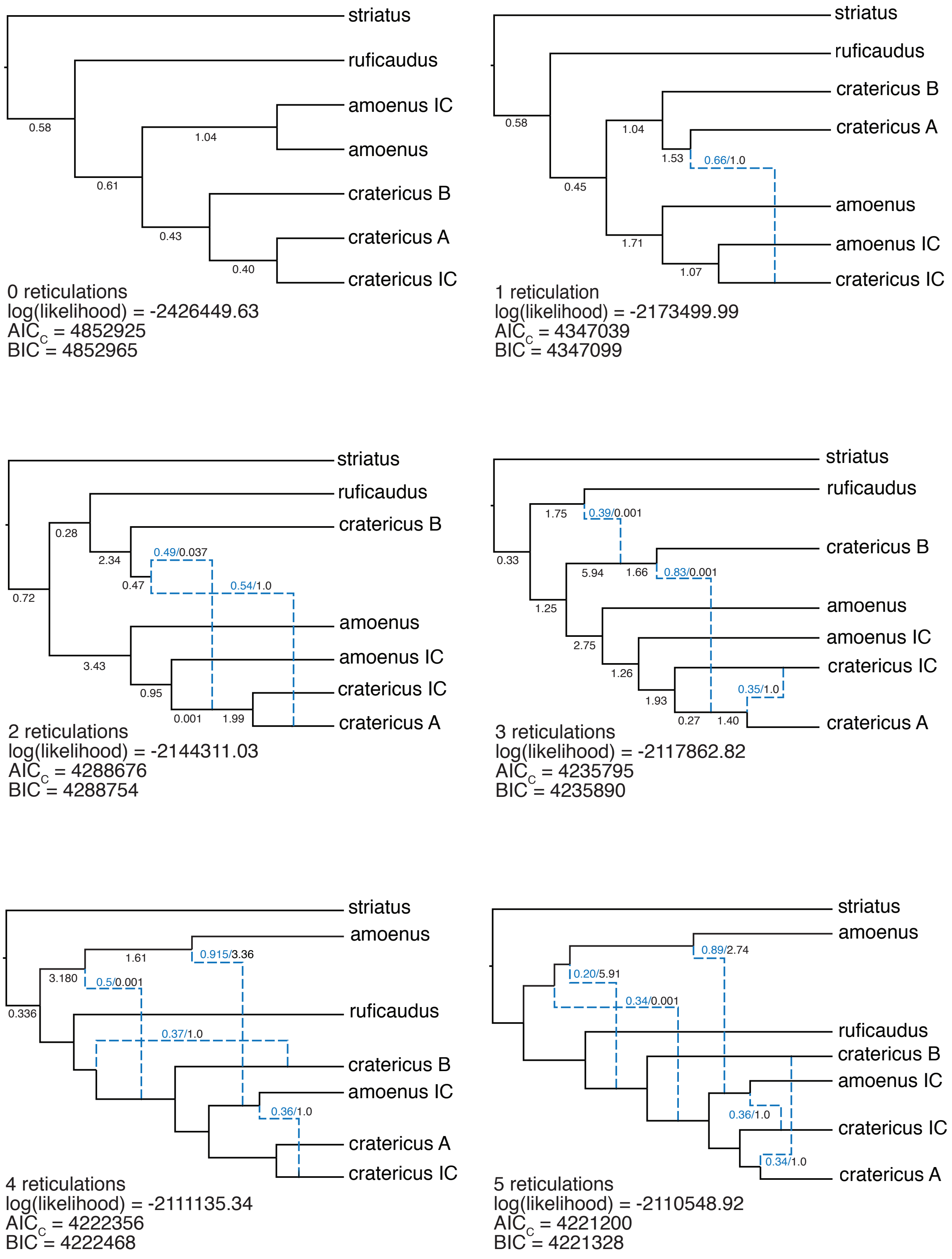


Figure S15 –Rooted pseudo-maximum likelihood networks for our expanded subset for *cratericus*, with 0 to 5 reticulations inferred from 161 gene trees with PhyloNet. Blue branches represent reticulations, branch lengths are represented in black and inheritance probabilities are represented in blue. Bellow each network, we report the log likelihood, Bayesian information criteria (BIC) and Akaike information criteria corrected for small sample sizes (AICc).

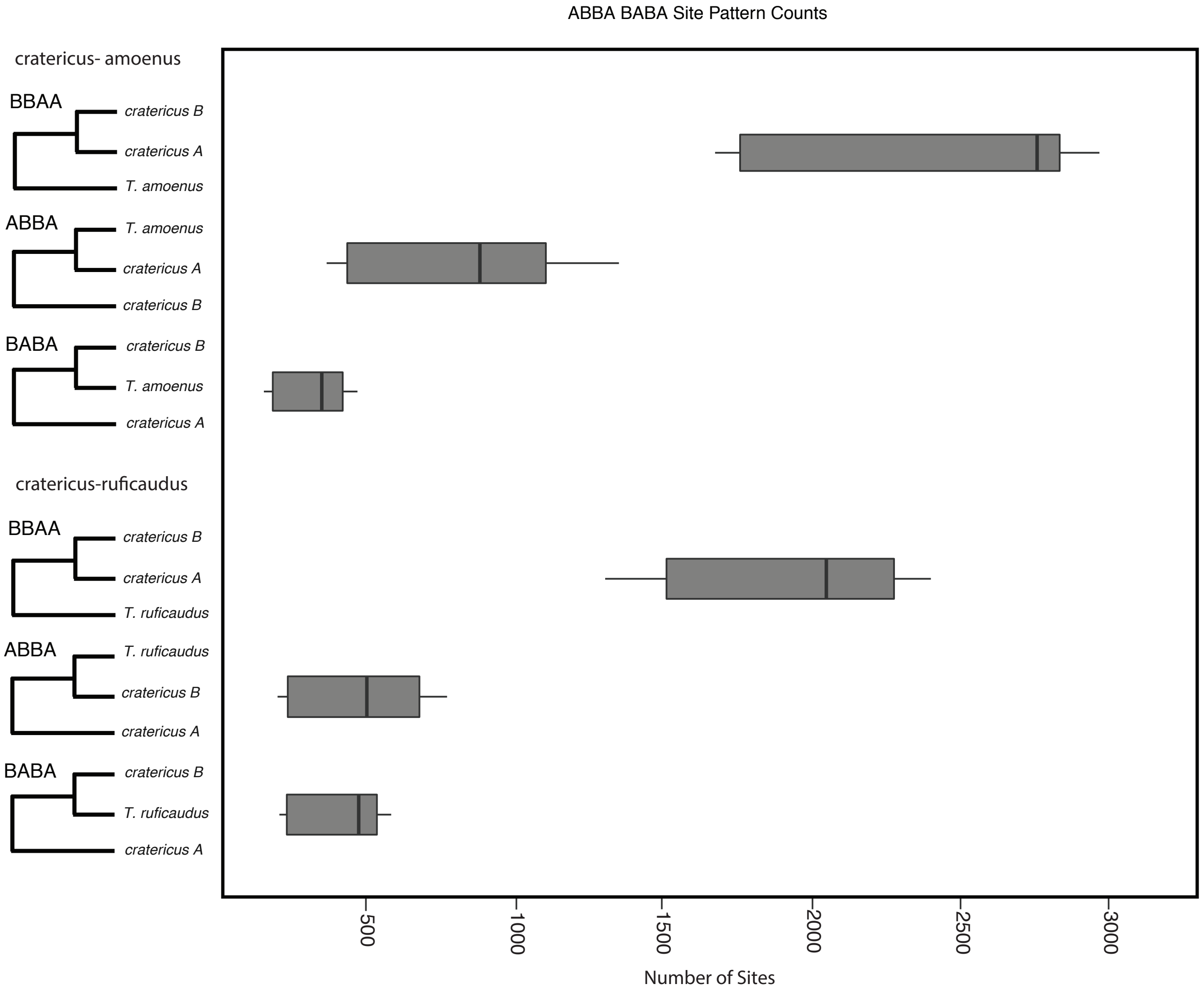


Figure S17 – Proportion of sites supporting the three possible rooted phylogenies between the two *cratericus* lineages and *T. amoenus* (top), and *T. ruficaudus* (bottom). Both contrasts show unequal representation of the two minor phylogenies and this evidence for introgression between *cratericus* lineage A and both *T. amoenus* (top) and *T. ruficaudus* (bottom).
